# PIEZO acts in an intestinal valve to regulate swallowing in *C. elegans*

**DOI:** 10.1101/2024.05.14.594054

**Authors:** Yeon-Ji Park, Jihye Yeon, Jihye Cho, Do-Young Kim, Xiaofei Bai, Yuna Oh, Jimin Kim, HoJin Nam, Hyeonjeong Hwang, Woojung Heo, Jinmahn Kim, Seoyoung Jun, Kyungeun Lee, KyeongJin Kang, Kyuhyung Kim

## Abstract

Sensations of the internal state of the body play crucial roles in regulating the physiological processes and maintaining homeostasis of an organism. However, our understanding of how internal signals are sensed, processed, and integrated to generate appropriate biological responses remains limited. Here, we show that the *C. elegans* PIEZO channel, encoded by *pezo-1*, regulates food movement in the intestine by detecting food accumulation in the anterior part of the intestinal lumen, thereby triggering rhythmical movement of the pharynx, referred to as the pharyngeal plunge. *pezo-1* deletion mutants exhibit defects in the pharyngeal plunge, which is rescued by PEZO-1 or mouse PIEZO1 expression, but not by PIEZO2, in a single isolated non-neuronal tissue of the digestive tract, the pharyngeal-intestinal valve. Genetic ablation or optogenetic activation of this valve inhibits or induces the pharyngeal plunge, respectively. Moreover, pressure built in the anterior lumen of the intestine results in a *pezo-1*-dependent pharyngeal plunge, which is driven by head muscle contraction. These findings illustrate how interoceptive processes in a digestive organ regulate swallowing through the PIEZO channel, providing insights into how interoception coordinates ingestive processes in higher animals, including humans.

## INTRODUCTION

Animals must detect and process external (exteroceptive) and internal (interoceptive) mechanical or physical stimuli in a process known as mechanotransduction for their survival and well-being ^1,2^. While specialized sensory organs detect mechanical forces to mediate touch, hearing, and proprioception, additional tissues and organs respond to internal mechanical cues ^3,4^. For instance, fluid flow and hydrostatic pressure influence the proper functions of tubular organs such as blood vessels, the urinary bladder, and the intestine. Dysregulation of these transduction processes results not only in sensory and motor defects but also in the disturbed homeostasis of development and physiology in various tissues and organs, leading to numerous disease statuses ^5^. However, the extent to which interoceptive mechanotransduction is involved in organ function is not fully understood.

Mechanosensory transduction pathways converge on distinct membrane proteins, including mechanosensitive ion channels, which convert mechanical stimuli into electrical or chemical signals depending on cell types ^6,7^. PIEZO ion channels, non-selective cationic channels directly activated by mechanical forces and evolutionarily well-conserved, have been shown to mediate mechanotransduction in various developmental and physiological contexts ^8^. The mammalian genes *Piezo1* and *Piezo2*, expressed in a wide range of tissues, including excitable and non-excitable tissues, play roles in diverse physiological processes, including circulation ^9–11^, touch sensation ^12^, proprioception ^13^, respiration ^14^, urination ^15^ by detecting blood flow, light-touch, muscle stretch, lung inflation, and bladder filling. Recent reports have shown that PIEZOs are widely expressed in the digestive system, including the gastrointestinal (GI) tract, and modulate gut motility and homeostasis, resulting in proper ingestion and digestion ^16–18^. Considering the broad expression of PIEZO channels in the digestive system, the precise roles of PIEZO channels in other digestive organs and their molecular mechanisms need to be further explored.

*C. elegans* exhibits a relatively simple digestive tract consisting of distinct anatomical and functional organs, including the mouth, pharynx, and intestine, allowing efficient and elaborate feeding and digestion of bacterial food ^19^. Ingested bacteria are ground and transported by pharyngeal pumping and peristalsis to the intestine, where further digestion and nutrient absorption occur ^20^. An additional valve-like organ or structure in the digestive tract, referred to as the pharyngeal-intestinal valve (PI valve), physically connects the pharynx to the intestine ^21^. This structure is anatomically analogous to the esophagus in vertebrates and appears to act as a barrier to control food flow ^22^. However, the precise roles of the PI valve in the *C. elegans* digestive system have not been explored. Furthermore, it is intriguing to decipher whether interoception is involved in the digestive system of *C. elegans* and, if so, which molecular sensors play a role. Here we find that the PI valve detects intestinal distention caused by food accumulation via *pezo-1*, a *C. elegans* ortholog of PIEZO channels, and triggers pharyngeal plunge to move food down to the intestine.

## RESULTS

### The *C. elegans* PIEZO channel, PEZO-1, mediates food movement in the intestine

The PIEZO genes are evolutionarily conserved across a wide range of organisms, including *C. elegans,* of which genome encodes one ortholog, *pezo-1* (Supplementary Fig. 1a). To investigate the function of *C. elegans* PEZO-1, we first sorted fourteen putative *pezo-1* isoforms into four groups based on the location of the translational start codon, which may share promoter regions, referred as p1, p2, p3, and p4 promoters, and observed their expression patterns (Fig. 1a). Consistent with previous reports ^23,24^, each promoter appeared to be expressed in a distinct type of cells; for example, the p1 promoter is specifically expressed in the pharyngeal-intestinal valve (PI valve), while the p3 promoter shows expression in several neurons and intestine (Fig. 1b, Supplementary Fig. 1b, and 1c). These findings suggest that similar to PIEZO genes in mammals, *C. elegans pezo-1* is expressed in various cell types throughout development, indicating diverse roles of *pezo-1* in *C. elegans* biology.

**Fig. 1:**
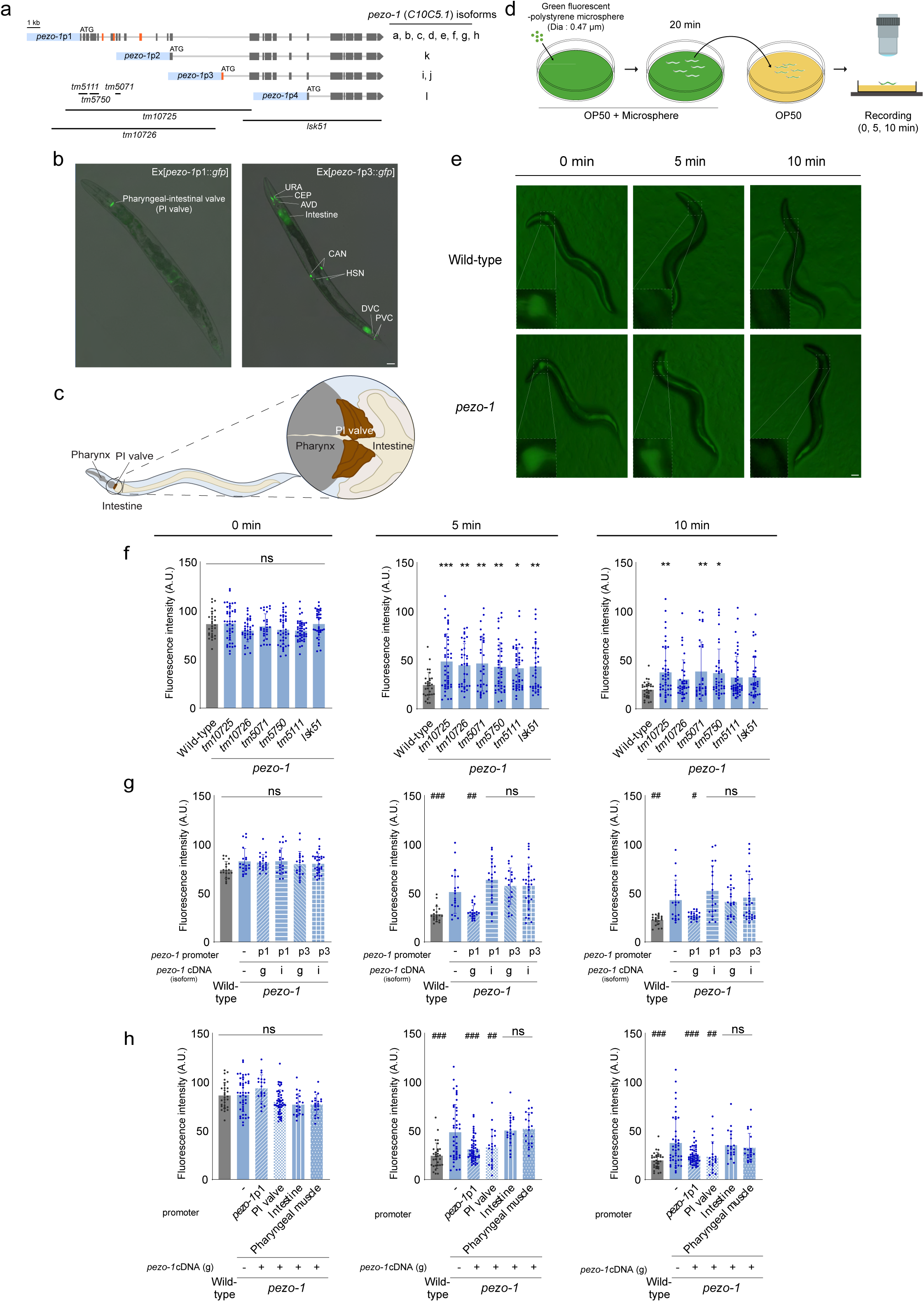
PEZO-1 regulates food movement in the intestine. (a) Genomic structure of the *pezo-1* gene. Orange bars and blue boxes represent alternatively spliced exons and promoter regions of *pezo-1* isoforms fused to the *gfp* gene, respectively. Black lines indicate deletion regions of *pezo-1* mutant alleles. (b) Representative images of wild-type animals expressing *pezo-1*p1::*gfp* (left) and *pezo-1*p3::*gfp* (right) transgene. Anterior is to the left. Scale bar: 10 μm. (c) Schematic of the digestive tract, including the pharynx, PI valve, and intestine in *C. elegans*. (d) Experimental scheme of the microsphere movement assay. Animals were placed in a plate containing green fluorescent-polystyrene microspheres seeded with bacterial food OP50 for 20 min, then transferred in an OP50-only plate during fluorescent imaging. (e) Representative images of the whole body of wild-type animal and *pezo-1* mutant taken at 0 min, 5 min, and 10 min after recovery from microsphere ingestion. Images in the lower left boxed regions are higher magnification of the boxed area (anterior part of the intestine). Scale bar: 10 μm. (f-h) Fluorescence intensity of microspheres in the anterior part of the intestine of (f) adult wild-type animals or *pezo-1* mutant alleles and (g-h) *pezo-1* (*tm10725*) rescued lines. Promoters and *pezo-1* cDNA isoforms used for rescue experiments are indicated. Each dot represents the fluorescence intensity of a single animal. n ≥ 20 for each genotype. Error bars indicate SD. *, **, ***, and #, ##, ### indicate significant differences from wild-type and *pezo-1* (*tm10725*) at p<0.05, p<0.01, and p<0.001, respectively (one-way ANOVA test followed by the Dunnett test).

Then we decided to uncover the *pezo-1* function in the PI valve, which consists of six epithelial cells and links the posterior of the pharynx to the anterior of the intestine ^22^ (Fig. 1c). Due to the lack of muscle fibers inside valve cells and synaptic innervation from the nervous system ^21^, it has been postulated that the valve could passively mediate food movement from the pharynx to the intestine, allowing further investigation of the PI valve’s function.

The expression of *pezo-1* in the PI valve prompted us to examine the flow of ingested food from the pharynx to the intestine. To track and monitor bacterial food movement, we fed animals with OP50-sized green fluorescent-polystyrene microsphere, the uptake of which could be comparable to OP50 ingestion ^25^ (Fig. 1d). In wild-type animals, microspheres ingested for 20 minutes moved from the mouth to the intestine but often accumulated in the anterior intestinal lumen near the PI valve (Fig. 1e, 1f, and Supplementary Fig. 1d). These residual microspheres in the anterior lumen gradually moved down the intestine and were eventually expelled by defecation in approximately 10 minutes when animals were fed with bacterial food ^25^ (Fig. 1e, 1f, and Supplementary Fig. 1e).

We then generated a *pezo-1* deletion mutant allele (*lsk51*) by using the CRISPR-Cas9 system, which deletes the C-terminus of all isoforms and thus could be a null allele (Fig. 1a) and observed the flow of microsphere in this mutant allele along with other available deletion alleles (Fig. 1a and 1d). While *pezo-1* mutants appeared to uptake microspheres normally, which then traveled into the anterior part of the intestine similar to wild-type animals, the intestinal accumulation of microspheres was more prominent in *pezo-1* mutants than in wild-type animals in 5 minutes or 10 minutes to a lesser extent (Fig. 1e, 1f, and Supplementary Fig. 1f). This indicates that in *pezo-1* mutants, ingested microspheres reside longer in the anterior intestinal lumen and move slowly down the intestinal lumen. Defecation-defective *unc-16* mutants, resulting from a lack of anterior body muscle contraction ^26^, were also defective in microsphere movement (Supplementary Fig. 2a), but *pezo-1* mutants were relatively normal in defecation (Supplementary Fig. 2b), suggesting that the *pezo-1* phenotype is not simply due to defecation defect.

Since we detected two abundant *pezo-1* transcripts through the reverse-transcription PCR, including *C10C5.1g* and *C10C5.1i* (Fig. 1a), rescue experiments were performed using these two cDNAs. The expression of a longer *pezo-1* cDNA (*C10C5.1g*), but not a shorter one (*C10C5.1i*), under the control of *pezo-1*p1 promoter rescued the defects of *pezo-1* mutants (Fig. 1g), indicating that full-length PEZO-1 is responsible for microsphere movement phenotype. Additionally, the PI valve-specific expression of a longer *pezo-1* cDNA fully rescued *pezo-1* defects, whereas pharyngeal muscle- and intestine-specific expression could not (Fig. 1h), suggesting that *pezo-1* acts in the PI valve to regulate microsphere movement in the intestine.

### PEZO-1 regulates pharyngeal plunge

We next sought to investigate the mechanisms by which accumulated microspheres move down the intestine independently of defecation. Interestingly, when worms were fed microspheres or even normal food, we observed the irregular but remarkably noticeable movement of the pharynx and PI valve, which together were pushed posteriorly and returned to their original position in a rhythmic fashion (Fig. 2a, 2b, and Supplementary Video 1). To quantify this phenotype, we measured the distance from the posterior end of the pharynx to the anterior end of the intestine, referred to as plunge length (Fig. 2a). When the pharynx of an adult worm moved into the intestine and the plunge length was greater than 3 μm (see Methods), we named this movement trait as a pharyngeal plunge (Fig. 2a and 2b). While wild-type animals triggered a pharyngeal plunge roughly every 4 seconds, and the average of their pharyngeal length was about 5 μm, *pezo-1* mutants exhibited strong defects in the pharyngeal plunge, with both pharyngeal frequency and length decreased (Fig. 2b-2d, Supplementary Fig. 2c, and Supplementary Video 2). Compared to defects in microsphere movement, *unc-16* mutants exhibited a normal pharyngeal plunge (Fig. 2d), further supporting the idea that defects in microsphere movement in *pezo-1* mutants are not due to defecation dysfunctions but rather defects in pharyngeal plunge.

**Fig. 2:**
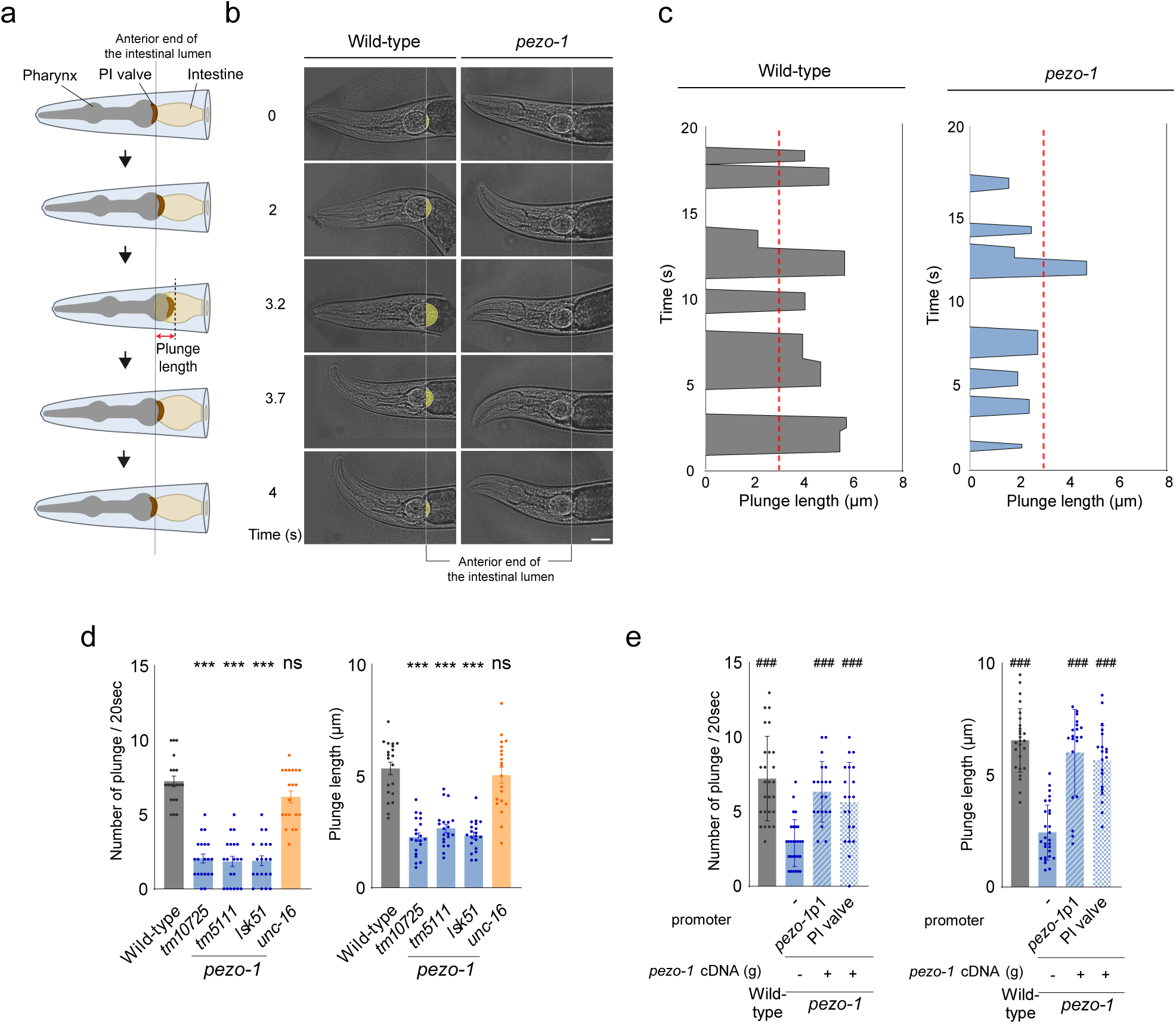
PEZO-1 mediates pharyngeal plunge. (a) Schematic representation of the pharyngeal plunge. Shown is the anterior digestive tract of a worm, comprising the mouth, pharynx, pharyngeal intestinal valve (PI valve), and intestine. Note the movement of the pharynx over time. (b) Representative images of worm’s head during the pharyngeal plunge over time in wild-type (left) and *pezo-1* mutant (right) animals over time. The white circle and vertical line indicate the terminal bulb of the pharynx and the anterior end of the intestine, respectively. Scale bar: 10 μm. (c) The time series plot of plunge length in wild-type (left) and *pezo-1* mutant (right) animals during 20 seconds pharyngeal plunge. A pharyngeal plunge is counted when the plunge length exceeds the red dotted line (3 μm). (d-e) Plunge frequency and plunge length of (d) wild-type animals and *pezo-1* mutant alleles and (e) *pezo-1* (*tm10725*) rescued lines. n ≥ 20 (plunge frequency) and N ≥ 20 (plunge length, three times per N) for each genotype. Error bars indicate SD for plunge frequency and SEM for plunge length. *** and ### indicate significant differences from wild-type and *pezo-1* (*tm10725*) at p<0.001, respectively. (one-way ANOVA test followed by the Dunnett test).

Consistent with microsphere movement, defects in the pharyngeal plunge of *pezo-1* mutants were fully rescued by the expression of a *pezo-1* cDNA (*C10C5.1g*) under the control of *pezo-1*p1 or the PI valve-specific promoter (Fig. 2e), providing further evidence that PEZO-1 acts in the PI valve to regulate pharyngeal plunge and thus food movement.

### Mouse *Piezo1* rescues defects in the pharyngeal plunge and microsphere movement of *pezo-1* mutants

PEZO-1 exhibits a high degree of homology in protein sequence and topology with mammalian PIEZO1 and PIEZO2, suggesting a functional conservation between these proteins^23,27^ (Fig. 3a). To investigate whether mammalian PIEZO channels can play comparable roles to the *C. elegans* PEZO-1, we expressed mouse *Piezo1* or *Piezo2* cDNA in the PI valve of *pezo-1* mutants and examined microsphere movement and pharyngeal plunge. The PI valve-specific expression of mouse *Piezo1,* but not *Piezo2,* fully rescued both defects in *pezo-1* mutants (Fig. 3b-3e), indicating the evolutionary conservation of PIEZO1 channel function in the PI valve between *C. elegans* and mice.

**Fig. 3:**
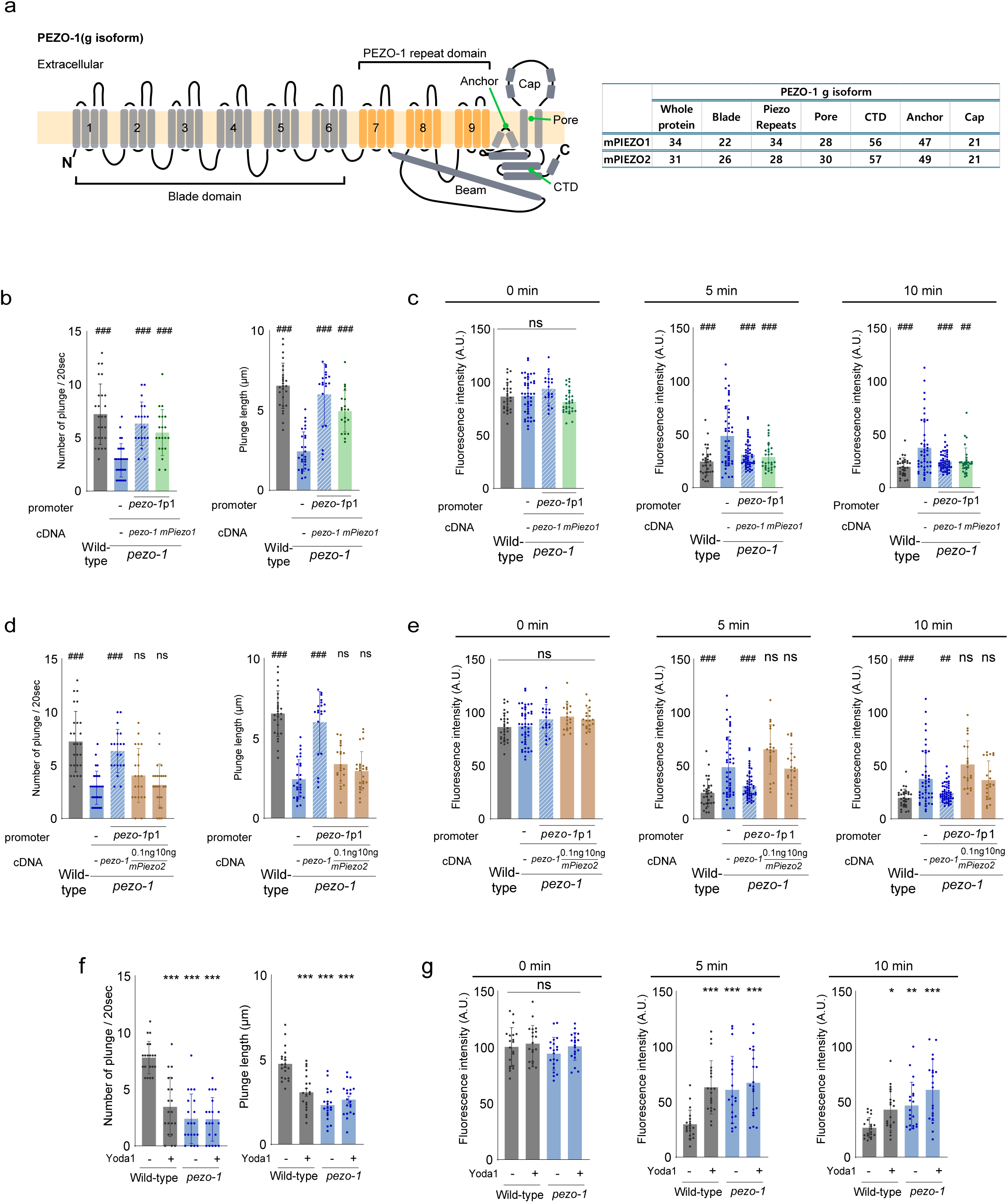
Expression of mouse *Piezo1* in the PI valve rescues defects of *pezo-1* mutants. (a) The architecture and topology of the PEZO-1 channel, along with the percentage similarity of amino acids between PEZO-1 and mPIEZO1 or mPIEZO2 in the whole protein and several functional domains (CTD : C-terminal domain). (b-e) Plunge frequency and plunge length (b, d), and fluorescence intensity of microspheres in the anterior part of the intestine (c, e) in wild-type animals, *pezo-1* (*tm10725*) mutants, and transgenic animals expressing either mouse *Piezo1* cDNA or mouse *Piezo2* cDNA in the PI valve. n ≥ 20 for each genotype. Error bars indicate SD for plunge frequency (b, d) and fluorescence intensity (c, e) and SEM for plunge length (b, d). ns, ##, ### indicate significant differences from *pezo-1* (*tm10725*) at no significant differences, p<0.01, and p<0.001, respectively. (one-way ANOVA test followed by the Dunnett test). (f) Plunge frequency and plunge length, and (g) fluorescence intensity of microspheres of wild-type and *pezo-1* mutant animals in the presence or the absence of 20 μM Yoda1. n ≥ 20 for each genotype. Error bars indicate SD. *, **, and *** indicate significant differences from wild-type in absence of Yoda1 at p<0.05, p<0.01, and p<0.001, respectively (one-way ANOVA test followed by the Dunnett test).

Yoda1 has been shown to open mammalian PIEZO1 channel, acting as a chemical agonist^28^, while the long-term effects of Yoda1 exposure on *C. elegans* appeared to inactivate PEZO-1 or PEZO-1-expressing cells ^23^. We found that wild-type animals fed with Yoda1 exhibited defects in pharyngeal plunge and microsphere movement similar to *pezo-1* mutants, whose defects were not aggravated by Yoda1 (Fig. 3f, 3g, and Supplementary Fig. 3a), further indicating that *pezo-1* indeed regulates pharyngeal plunge and food movement and *mPiezo1* exhibit similar functions to *pezo-1*.

### The PI valve mediates pharyngeal plunge, which forces food to move down the intestine

Identification of the PI valve as the site of action of *pezo-1* allowed us to explore the roles of the PI valve in pharyngeal plunge and microsphere movement. We decided to optogenetically ablate the PI valve by expressing the genetically encoded photosensitizer miniSOG (mini Singlet Oxygen Generator) ^29,30^. Upon exposure to blue light, transgenic animals expressing *miniSOG* and *gfp* (green fluorescent protein) genes in the PI valve appeared to lose *gfp* expression in an hour (Supplementary Fig. 3b), likely due to the cell death, and showed decreased pharyngeal plunge and microsphere movement comparable to *pezo-1* mutants (Fig. 4a and 4b), indicating that the PI valve is required for pharyngeal plunge and microsphere movement.

**Fig. 4:**
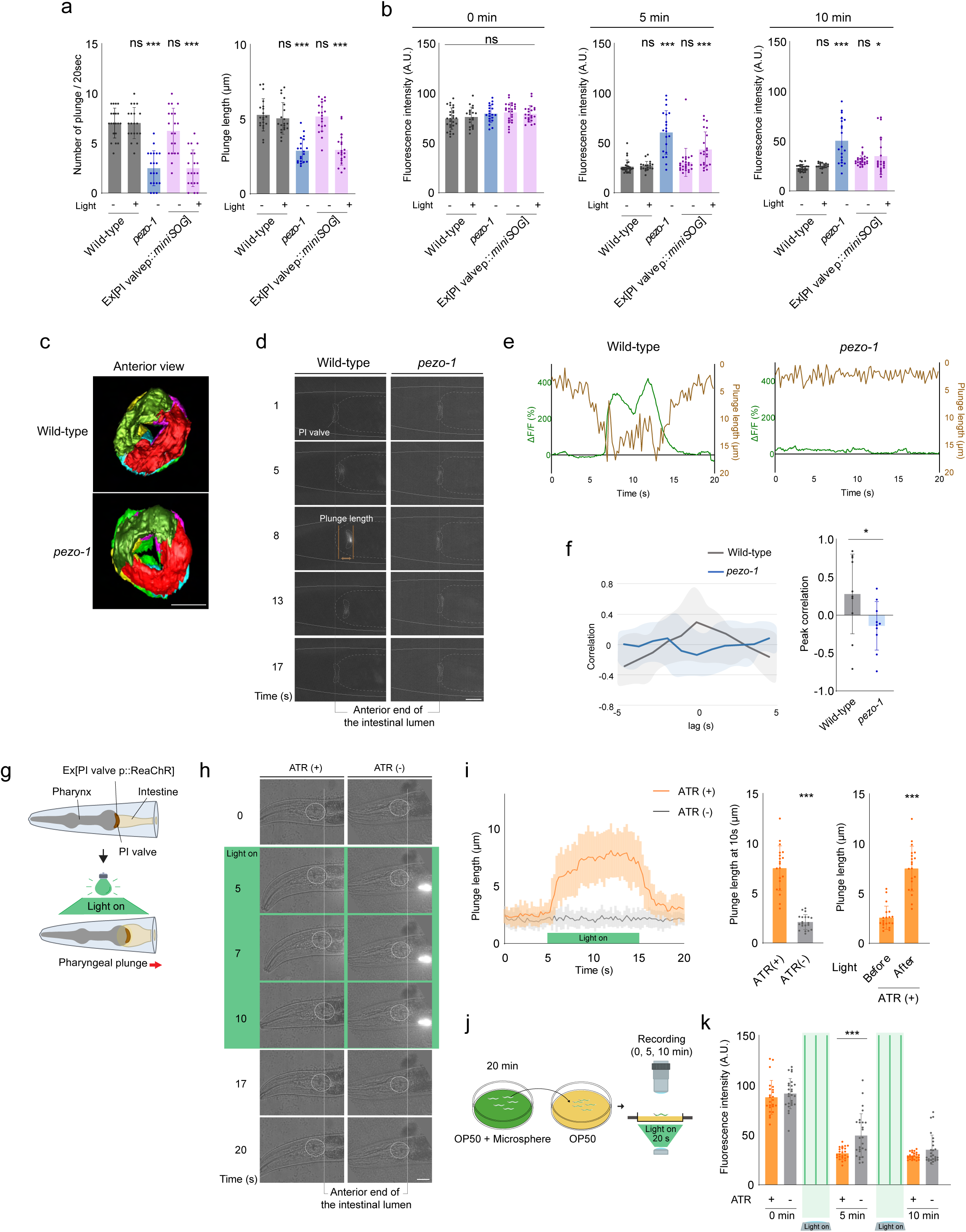
The PI valve elicits pharyngeal plunge, which facilitates food movement in the intestine. (a-b) Plunge frequency and plunge length (a), and fluorescence intensity of microspheres in the anterior part of the intestine (b) in wild-type animals, *pezo-1* (*tm10725*) mutants, and transgenic animals expressing the *miniSOG* gene in the PI valve. n ≥ 20 for each genotype. Error bars indicate SD for plunge frequency (a) and fluorescence intensity (b) and SEM for plunge length (a). * and *** indicate significant differences from light-unexposed wild-type at p<0.05 and p<0.001 (one-way ANOVA test followed by the Dunnett test). (c) Serial Block Face Scanning Electron Microscopy images of the anterior view of the PI valve in wild-type animals and *pezo-1* (*tm10725*) mutants. Each of the PI valve cells is color-labeled in green (vpi3), red (vpi3), yellow (vpi2), magenta (vpi2), cyan (vpi2) and light green (vpi1) respectively. Scale bar: 5 μm. (d) Representative images of the worm’s head during the pharyngeal plunge over time in wild-type and *pezo-1* (*tm10725*) mutant animals expressing GCaMP6s in the PI valve. The white circle and vertical line indicate the PI valve and the anterior end of the intestine, respectively. The distance between brown lines indicates the plunge length. Scale bar: 10 μm. (e) Representative calcium dynamics (green) in the PI valve and the corresponding plunge length (brown) in the same wild-type or *pezo-1* (*tm10725*) mutant animals. (f) Cross-correlations (left) and peak correlations (right) between PI valve calcium responses and Pharyngeal plunge length. The Peak correlation value is obtained from lag 0 of cross-correlation. *n* = 15 for each. Error bars indicate SD. * indicates significant differences from wild-type at p < 0.05 (unpaired t-test). (g) Experimental scheme of optogenetic activation of the PI valve by expressing ReaChR under the control of PI valve-specific *valv-1* promoter. (h) Representative images of a *valv-1*p::ReaChR::mKate2 transgenic animal during light stimulation over time. Each animal is exposed to light stimulation in the presence or absence of all trans retinal (ATR). The white circle indicates the terminal bulb of the pharynx and the white line indicates the anterior end of the intestinal lumen. Scale bar: 10 μm. (i) The time series plot of plunge length of *valv-1*p::ReaChR::mKate2 transgenic animals during light stimulation (left). Plunge length after 10 seconds of light stimulation in the presence or absence of ATR (middle) and before and after light stimulation in the presence of ATR (right). (j) Experimental scheme for the microsphere movement assay upon optogenetic activation of the PI valve. Animals are exposed to three 20-seconds light pulses at 2-3 and 7-8 minutes. (k) Fluorescence intensity of microsphere in the anterior part of the intestine of *valv-1*p::ReaChR::mKate2 transgenic animals. Each dot represents the fluorescence intensity of a single animal. n ≥ 20 each. Error bars indicate SD. *** indicate significant differences at p<0.001 (unpaired *t*-test).

We then examined whether the structure of the PI valve is abnormal in *pezo-1* mutants and observed the morphology of the PI valve by Serial Block Face Scanning Electron Microscopy (SBF-SEM) ^31^. The overall morphology and internal structures did not appear to be altered in *pezo-1* mutants (Fig. 4c and Supplementary Fig. 3c), suggesting that *pezo-1* mutant phenotypes are not simply due to structural defects of the PI valve. Furthermore, *pezo-1* mutations did not affect the expression of the PI valve-expressed genes, including LIM zinc-binding domain-containing protein VALV-1 ^32^ and transmembrane protein MIG-13 ^33^ (Supplementary Fig. 4a), indicating that overall gene expression in the PI valve may not be affected in *pezo-1* mutants.

To further probe the roles of the PI valve, we expressed the genetically encoded calcium sensor GCaMP6s ^34^ in the PI valve and monitored, if any, calcium transients as the pharyngeal plunge occurs. We observed a transient increase in the PI valve Ca^2+^ levels when the pharyngeal plunge was triggered (Fig. 4d, 4e, and Supplementary Fig. 4b). PI valve Ca^2+^ transients were returned to the baseline as the pharyngeal plunge was terminated (Fig. 4d, 4e, and Supplementary Video 3). Cross-correlation analysis of the PI valve Ca^2+^ transients and pharyngeal plunge revealed a strong correlation between the PI valve Ca^2+^ activity and the pharyngeal plunge (Fig. 4f). These Ca^2+^ transients were abolished in *pezo-1* mutants (Fig. 4d, 4e, Supplementary Fig. 4b, and Supplementary Video 4), suggesting that PEZO-1 facilitates the passage of Ca^2+^ into the PI valve, resulting in the pharyngeal plunge.

We next examined whether optogenetic excitation in the PI valve induces the pharyngeal plunge and thus microsphere movement by expressing the channel rhodopsin variant ReaChR ^35^ in the PI valve (Fig. 4g). Light stimulation in the presence of all-trans-retinal (ATR) instantly triggered the pharyngeal plunge of transgenic animals, which persisted until the light was turned off in 10 seconds (Fig. 4h, 4i, Supplementary Fig. 4c, and Supplementary Video 5, Video 6). Moreover, repeated 20-second pulses of light also increased microsphere movement (Fig. 4j and 4k), indicating that pharyngeal plunge indeed promotes microsphere movement. Together, these results indicate that the activities or Ca^2+^ levels in the PI valve modulate the pharyngeal plunge and, consequently, microsphere movement.

### PEZO-1 induces the pharyngeal plunge by detecting distension built in the anterior part of **the intestine**

To investigate the roles of PEZO-1 in the PI valve underlying the pharyngeal plunge, we examined the subcellular localization of PEZO-1 in the PI valve by endogenously tagging the C-terminal of PEZO-1 with mScarlet fluorescent protein ^23^. Consistent with promoter activities of the *pezo-1* gene, PEZO-1 was expressed in the PI valve and predominantly localized in the cell membrane facing the intestine but not the pharynx (Fig. 5a), suggesting that PEZO-1 detects mechanical force or pressure generated from the anterior part of the intestine in order to mediate the pharyngeal plunge.

**Fig. 5:**
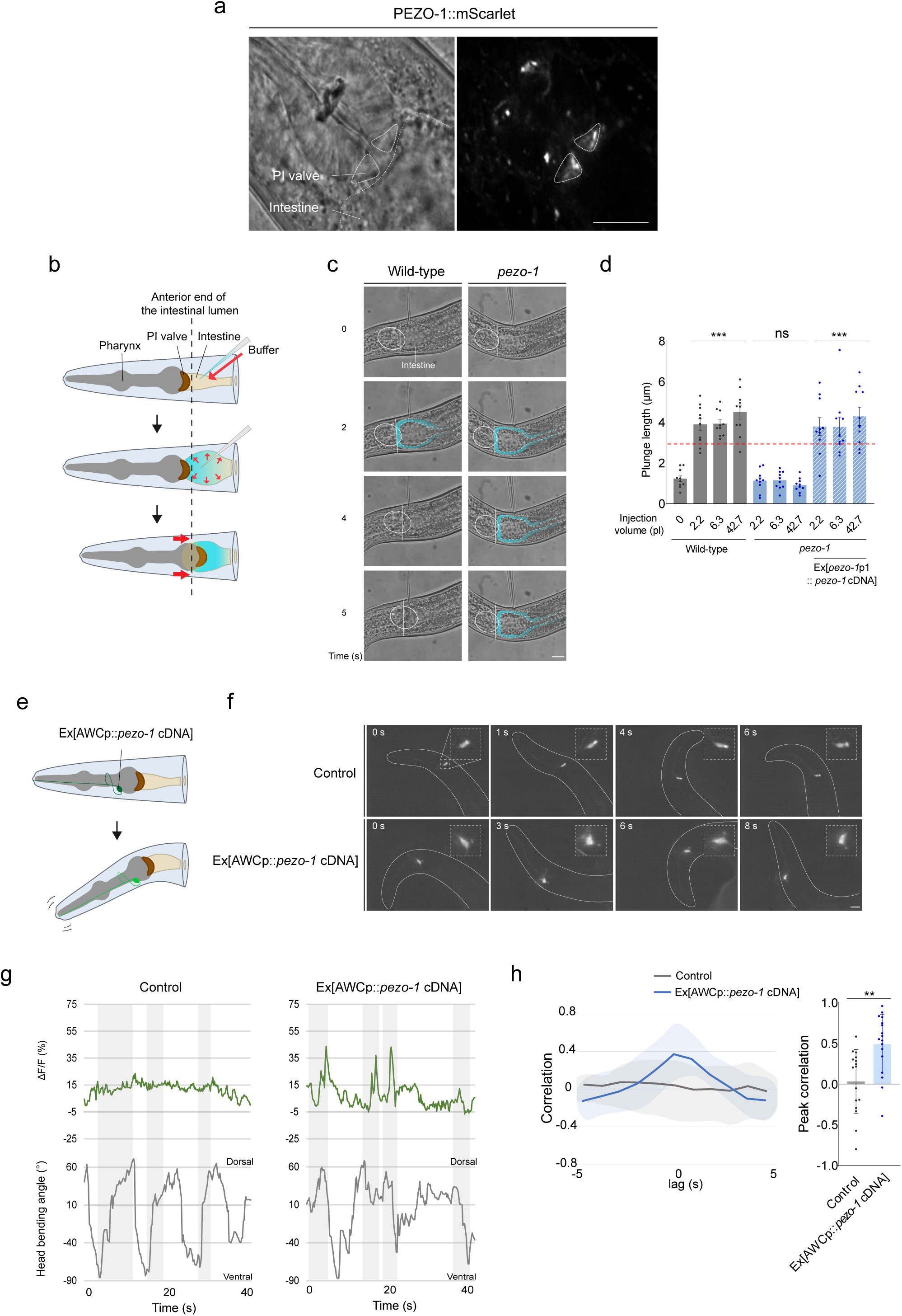
Distension built in the anterior part of the intestine triggers PEZO-1-mediated pharyngeal plunge. (a) Bright field (left) and fluorescent (right) images of transgenic animals expressing PEZO-1::mScarlet fusion proteins in the PI valve. Scale bar: 10 μm. (b) Experimental scheme for inflating the anterior part of the intestine. (c) Representative images of worm’s head during injecting buffer in wild-type (left) and *pezo-1* mutant (right) animals over time. The white circle and vertical line indicate the terminal bulb of the pharynx and the anterior end of the intestine, respectively. The blue-colored region indicates an inflated area in the intestine after buffer injection. Scale bar: 10 μm. (d) Plunge length in wild-type animals or *pezo-1* mutants after inflating the anterior part of the intestine. n= 20 for each genotype. Error bars indicate SD. *** indicates significant differences from wild-type without buffer injection (0 pl) at p<0.001 (one-way ANOVA test followed by the Dunnett test). (e) Experimental scheme for PEZO-1 ectopic expression in the AWC neurons. (f) Representative single-frame images of GCaMP3 signal in the AWC soma of freely moving control or a AWCp::*pezo-1* cDNA transgenic animals during head bending over time. Images in the upper right boxed regions are higher magnification of the AWC soma. Scale bar: 10 μm. (g) Representative trace of AWC GCaMP activities (upper) and the corresponding head bending angle (lower) of control (left) or an AWCp::*pezo-1* cDNA transgenic (right) animal. (h) The cross-correlation curve (left) and peak correlation (right) between AWC calcium responses and head bending. n = 15. Error bars indicate SD, and *** indicates significant differences from control at p<0.001 (unpaired t-test).

Based on the compartmentalized localization of the PEZO-1 on the surface of PI valve and the acute accumulation of food in the anterior part of the intestine, we hypothesized that the PI valve may respond to pressure or distension built in the anterior part of the intestine resulting from food accumulation via the PEZO-1. To test this hypothesis, we inflated the anterior part of the intestine by microinjection of buffer solution (Fig. 5b). Microinjection of approximately 2-50 pl buffer induced expansion of the anterior intestinal lumen (Fig. 5c and Supplementary Fig. 5a), and remarkably triggered pharyngeal plunge in wild-type animals and rescued lines in 2-3 seconds after injection, but not in *pezo-1* mutants (Fig 5c, 5d, and Supplementary Video 7, Video 8). These results indicate that pharyngeal plunge can be elicited by distension of the intestine depending on PEZO-1.

A previous study has shown that PEZO-1 could be gated by mechanical pressure in heterologous systems ^24,36^. To further support the mechanosensitive function of PEZO-1, we exploited an alternative approach ^37^ in which PEZO-1 is ectopically expressed in the AWC chemosensory neurons of transgenic animals expressing GCaMP3 and monitored the AWC Ca^2+^ activities during freely moving condition (Fig. 5e). Consistent with previous observations ^37^, the AWC neurons of wild-type animals exhibited inconsistent and weak Ca^2+^ signals when they moved forward (Fig. 5f, 5g, and Supplementary Fig. 5b). However, transgenic animals expressing PEZO-1 in the AWC neurons showed reliable and strong Ca^2+^ transients that were correlated with head-bending (Fig. 5f-5h), indicating that ectopic expression of PEZO-1 is sufficient to confer neuronal responses in the AWC neurons upon head deformation. These results further suggest that PEZO-1 acts as a mechanosensitive receptor in the context of the PI valve-mediated pharyngeal plunge.

### The pharyngeal plunge is driven by head/neck muscle contraction via chemical synaptic transmission

We next sought to investigate the forces responsible for pharyngeal movement. Consistent with the previous report ^22^, we could not observe any muscle fibers inside the PI valve (Fig. 4c and Supplementary Fig. 3c), excluding the possibility of the PI valve contraction as a force source. Since the PI valve directly connects to a pm8 pharyngeal muscle via gap junctions ^21,22^, we examined the contraction and Ca^2+^ activities of the pharyngeal muscles during the pharyngeal plunge and found no noticeable activities of pharyngeal muscles (Supplementary Fig. 6), suggesting that pharyngeal plunge is not forced by pharyngeal muscle contraction. The anterior, but not other parts of, the pharynx is anchored to the buccal cavity or mouth and attached to head muscles via tendon-like structures ^38^ (Fig. 6a). Unlike pharyngeal muscles, head/neck muscles exhibited strong contraction and Ca^2+^ transient during the pharyngeal plunge (Fig. 6b and 6c), indicating that contraction of head/neck muscles forces the pharynx to move backward.

**Fig. 6:**
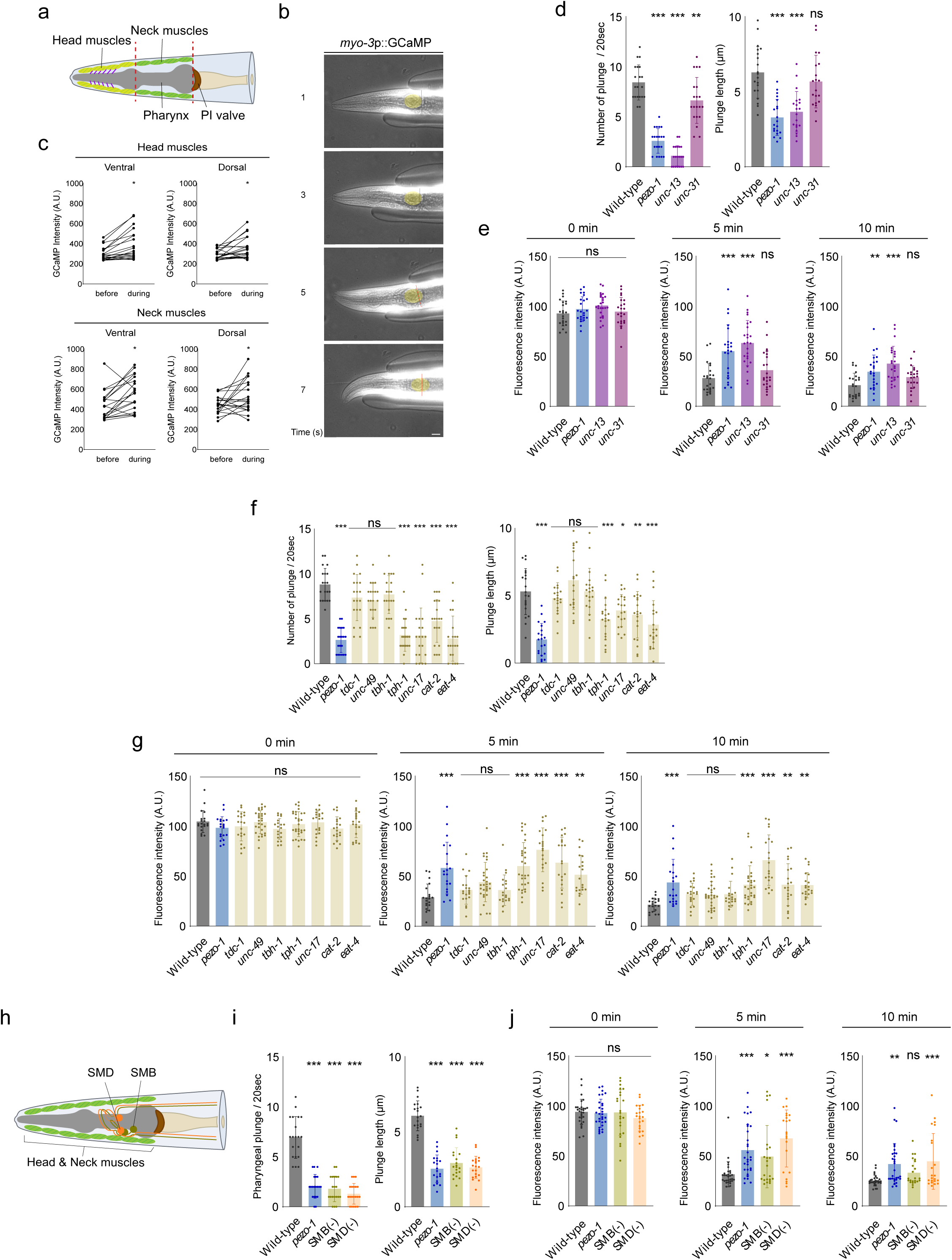
Head/neck muscles contraction mediates pharyngeal plunge. (a) Schematic of the head muscles (light green), neck muscles (green), and tendon-like structure (purple) in *C. elegans*. (b) Representative images of worm’s head during the pharyngeal plunge over time in the body wall muscles of wild-type animals expressing GCaMP3. The yellow circle and red line indicate the terminal bulb and the anterior end of the intestine, respectively. Scale bar: 10 μm. (c) The head and neck muscles activities before or during the pharyngeal plunge. n=20. Error bars indicate SD and * indicates significant differences at p<0.05 (unpaired t-test). (d-g) Plunge frequency and plunge length (d, f), and fluorescence intensity of microspheres in the anterior part of the intestine (e, g) in indicated genotypes. n ≥ 20 for each genotype. Error bars indicate SD for plunge frequency and fluorescence intensity and SEM for plunge length. *, **, and *** indicate significant differences from wild-type at p<0.05, p<0.01, and p<0.001 (one-way ANOVA test followed by the Dunnett test). (H) Schematic of the SMB (dark green) and SMD (orange) motor neurons in *C. elegans*. Dorsal left and ventral left cell bodies are shown. (i) Plunge frequency and plunge length, and (j) fluorescence intensity of microspheres in the anterior part of the intestine in indicated genotypes. n ≥ 20 for each genotype. Error bars indicate SD for plunge frequency and fluorescence intensity and SEM for plunge length. *, **, and *** indicate significant differences from wild-type at p<0.05, p<0.01, and p<0.001 (one-way ANOVA test followed by the Dunnett test).

Since head/neck muscle contraction could be elicited by chemical synaptic transmission and/or peptidergic signaling, we examined pharyngeal plunge and microsphere movement in UNC13/*unc-13* and CAPS/*unc-31* mutants, which are defective in synaptic vesicle release and dense core vesicle release, respectively ^39,40^. *unc-13*, but not *unc-31*, mutants exhibited strong defects in pharyngeal plunge and microsphere movement comparable to *pezo-1* mutants (Fig. 6d and 6e), suggesting that small molecule, but not large neuropeptide, chemical synaptic transmission mediates pharyngeal plunge. We further investigated which neurotransmitter mediates pharyngeal plunge and microsphere movement by examining neurotransmitter mutants, including *tdc-1* (tyramine), *unc-49* (GABA), *tbh-1* (octopamine), *tph-1* (serotonin), *unc-17* (acetylcholine), *cat-2* (dopamine) or *eat-4* (glutamate) ^41^, and found that *tph-1*, *unc-17, cat-2*, *eat-4,* but not *tdc-1*, *unc-49*, *tbh-1* mutants exhibited defects in both pharyngeal plunge and microsphere movement (Fig. 6f and 6g), indicating that multiple neurotransmitters are involved in these processes.

The SMB and SMD neurons are two major types of motor neurons that innervate the head and neck muscles ^22^ (Fig. 6h). We tested transgenic animals in which the SMB or SMD neurons are genetically ablated and found that these animals exhibited strong defects in pharyngeal plunge and microsphere movement comparable to *pezo-1* mutants (Fig. 6i and 6j), indicating that the SMB and SMD motor neurons play roles in mediating pharyngeal plunge. Together, the PI valve transmits the interoceptive signal from the intestine to head/neck muscles through multilayer synaptic transmission and motor neurons.

## DISCUSSION

*C. elegans* perceives and responds to external cues with remarkable sensitivities, as a form of exteroception, and specialized sensory cells and receptor molecules to detect exteroceptive stimuli underlying chemosensation, thermosensation, light sensation, and touch sensation have been identified ^42^. However, despite its well-established significance in higher organisms, including humans, it has been intriguing whether interoception, the internal sensing of physiological states, especially via mechanotransduction, play a crucial role in *C. elegans* biology. Here we demonstrate that *C. elegans* detects and integrates internal mechanical signals associated with the digestive process to facilitate food movement.

In *C. elegans*, the initial movement and processing of bacterial food in the digestive tract are mediated by the pharynx where two modes of the pharyngeal muscle contraction, referred as pumping and isthmus peristalsis, create a flow of partially digested bacteria within the pharynx^43^. The flow of liquid from the pharynx into the intestine is not driven by peristaltic contractions, as seen in more complex digestive systems, but the difference in timing between pumping and isthmus peristalsis seems to propel bacteria posteriorly to the intestine ^44,45^. Inside the intestine, bacterial food movement is regulated by a coordinated and rhythmic defecation process in which multiple layers of muscles play distinct roles. Specifically, posterior and anterior body wall muscles contract sequentially, squeezing bacterial food anteriorly and then posteriorly ^26^. Here we have presented several pieces of evidence indicating that the pharyngeal plunge further facilitates bacterial food movement in the intestine. First, *pezo-1* deletion mutants exhibit severe defects in the pharyngeal plunge as well as microsphere movement in the intestine that are rescued by the expression of PEZO-1 in the PI valve, but neither in the intestine or in the pharyngeal muscles. Second, optogenetic activation or inhibition of the PI valve results in increased or decreased pharyngeal plunge and microsphere movement, respectively. Third, the pharyngeal plunge occurs independently of defecation; although these processes follow a rhythmic pattern, defecation happens every 45 seconds whereas the pharyngeal plunge occurs roughly 4-5 seconds with less consistency. In addition, *unc-16* mutants, that exhibit severe defects in anterior body wall muscle contraction and thus defection, are normal in pharyngeal plunge while *pezo-1* mutants are relatively normal in defecation. Lastly, the pharyngeal plunge is also a separate event from pharyngeal pumping, as no calcium activities in pharyngeal muscles are detected during the pharyngeal plunge and optogenetic activation of the PI valve could not elicit pharyngeal muscle contraction. Together, while defecation could be a significant driving force in moving bacterial food down to the posterior intestine, we ensure that the pharyngeal plunge is a highly elaborate but delicate form of digestive processes exploited by the PI valve.

What physiological roles does PEZO-1 play in the PI valve to mediate the pharyngeal plunge? Our data from subcellular localization, Ca^2+^ imaging, optogenetics, and intestine-inflating experiments, support a model in which the nonselective cationic ion channel, PEZO-1, acts as a mechanosensitive receptor in the PI valve that detects mechanical pressure or distension built in the anterior part of the intestine. Upon mechanical activation, PEZO-1 facilitates the influx of Ca^2+^ ions into the PI valve, leading to Ca^2+^ transient, possibly initiating intracellular second messenger signaling pathway and thus triggering the pharyngeal plunge (Fig. 7).

**Fig. 7:**
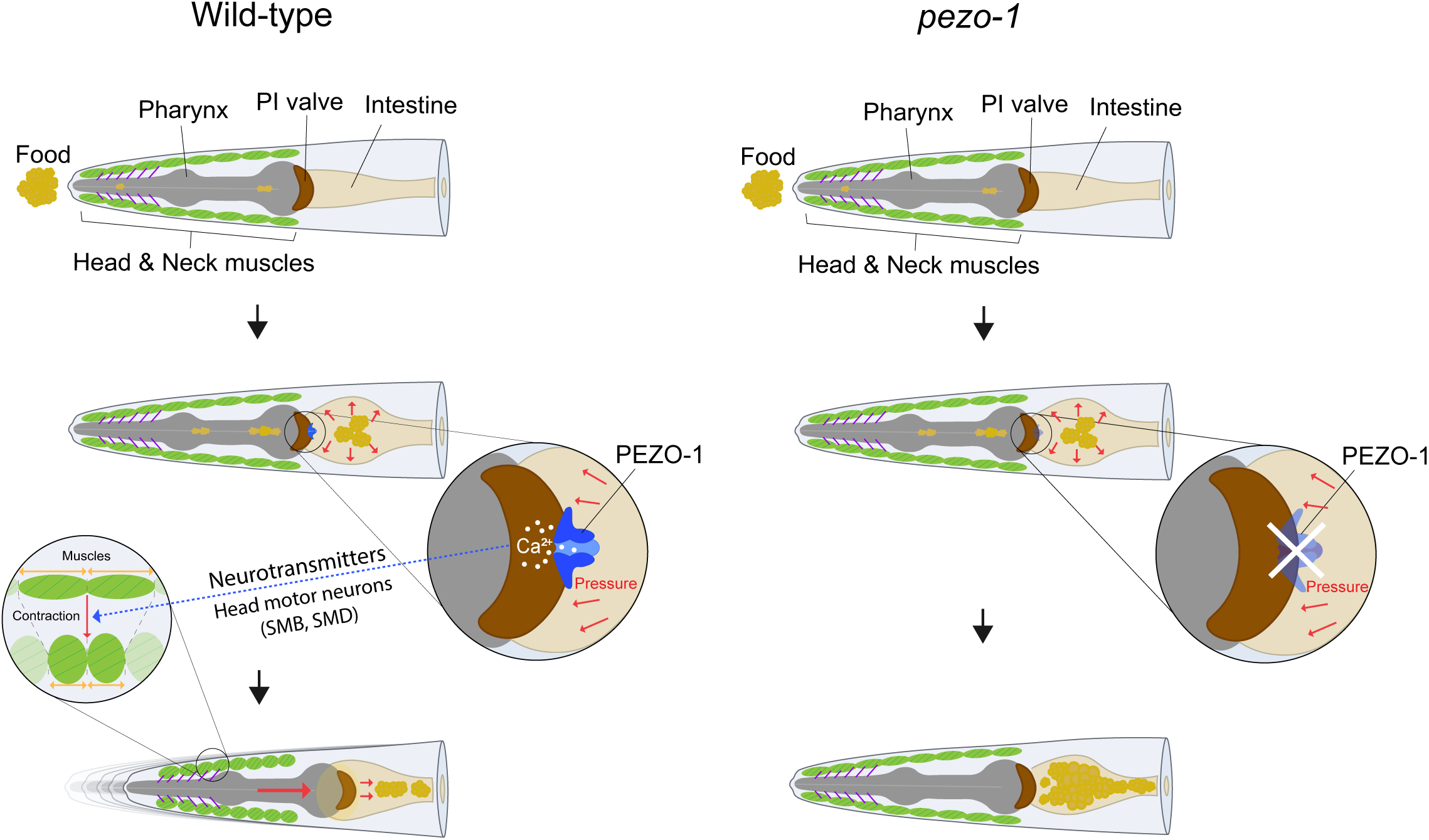
Model illustrating the functions of *pezo-1* in the PI valve-mediated pharyngeal plunge. *C. elegans* ingests food, which moves posteriorly and accumulates in the anterior part of the intestine. PEZO-1 channels expressed in the cell membrane of the PI valve, detect distension or pressure from the anterior intestinal lumen, resulting in calcium transients in the PI valve. Calcium activities in the PI valve elicit contraction of head/neck muscles via several neurotransmitters and head motor neurons, triggering pharyngeal plunge that pushes food into the posterior intestine. This process is illustrated in the images, where wild-type animals (left) show normal pharyngeal plunge, while *pezo-1* mutants (right) exhibit defects in pharyngeal plunge.

Compared to the predominant expression pattern of mammalian PIEZO1 in non-neuronal tissues and PIEZO2 in neuronal tissues, *C. elegans* PEZO-1 isoforms display either a non-neuronal or neuronal expression pattern; for example, *g* or *i* isoforms are expressed in the non-neuronal PI valve or distinct neuronal cell-types, respectively. Interestingly, the *g* isoform exhibits a stronger sequence similarity to the mouse PIEZO1 than PIEZO2 at the protein level ^23^. Although electrophysiological properties of the PEZO-1 *g* isoform recapitulate those of mammalian PIEZO channels, including voltage-dependent inactivation and nonselective cation currents, it is hard to determine whether the *g* isoform is more similar to PIEZO1, probably due to different heterologous systems used in previous studies ^8,24,46^. Our rescue data strongly indicates that mouse PIEZO1, but not PIEZO2, can functionally substitute for the PEZO-1 *g* isoform underlying the pharyngeal plunge. Moreover, PIEZO1 selective agonist, Yoda1, also exerts the pharyngeal plunge ^28^. It is intriguing to investigate whether the neuronally expressed PEZO-1 *i* isoform exhibits functional conversation with PIEZO2 for a future study.

The previous EM analysis, along with our own findings, indicates that the PI valve lacks muscle fibers and synaptic innervation from the nervous system ^22^. Given this, how do Ca^2+^ transients in the PI valve lead to the head/neck muscle contraction? Since the PI valve is directly connected to the pm8 pharyngeal muscle via a few gap junction proteins and the tendonous structure anchors the pharynx to head muscles ^38^, one possibility is that Ca^2+^ signals are transmitted to the head/neck muscles through the activation of pharyngeal muscles. However, our Ca^2+^ imaging and optogenetic results showing that pharyngeal muscle contraction is not involved in the pharyngeal plunge, suggest that this is not the case. Another possibility is that, similar to enterochromaffin cells in the mammalian small intestine and colon, where serotonin (5-HT) is secreted to neuronal and non-neuronal cells, influencing various physiological processes ^47^, the PI valve can release neurotransmitters as paracrine singling molecules to affect neighboring neuronal processes, including the SMB/SMD head motor neurons ^37,48^. Due to the multiple neurotransmitters involved in the pharyngeal plunge (Fig. 6), it would be intriguing to explore this possibility in the future. Together, our results not only contribute to our understanding of interoception in a simpler organism but also underscore the evolutionary conservation of key molecular pathways involved in the internal sensory perception across different species. Studying PEZO-1 mediated interoception in *C. elegans* provides insights into the internal sensing mechanisms within the context of gastrointestinal processes and also serves as an attractive *in vivo* drug screen platform to identify novel chemicals interacting with PIEZO channels of higher animals, including humans.

## METHODS

### C. elegans Strains

All strains were maintained at 20°C^49^. The N2 Bristol strain was used as a wild-type strain. The deletion sites of *pezo-1 (tm10725, tm10726, tm5071, tm5750,* and *tm5111)* were identified by PCR (*tm10725* Forward primer : GACGAGGAAGGATAGCGCAA, *tm10725* Reverse primer : AAGTGCGCGTCCAAAGACCA, *tm10726* Forward primer : AAAAGGCGGGTCCCTTTCAC, *tm10726* Reverse primer : CTAGAGCCCCAGAATCTAAC, *tm5071* Forward primer : CACCCCCAGCTCCAGTCTTT, *tm5071* Reverse primer : CATTACTCCTCCTACCCTGT, *tm5750* Forward primer : CGCCCGTAGTTAAAGGGGAT, *tm5750* Reverse primer : AATGACGGTCCCGCCGCTTT, *tm5111* Forward primer : CGCCCGTAGTTAAAGGGGAT, *tm5111* Reverse primer : AATGACGGTCCCGCCGCTTT). To generate the *pezo-1(lsk51)*, the CRISPR-Cas 9 system (*pezo-1* sg Forward primer : GTTAGCATATTACGATCCGCGTTTTAGAGCTAGAAAT, *pezo-1* sg Reverse primer : GCGGATCGTAATATGCTAACAAGACATCTCGCAATAG, *pezo-1* 3’ utr sg Forward primer : GTCCATTCGATGAGTGTCGCGTTTTAGAGCTAGAAAT, *pezo-1* 3’ utr sg Reverse primer : GCGACACTCATCGAATGGACAAGACATCTCGCAATAGG) was used and the genomic deletion of *pezo-1* was confirmed by PCR (*pezo-1* deletion right primer : TCAACTCGAACGACAAGCTG, *pezo-1* deletion left primer : CCCTGGTTGCCTTGTTTTAT). All the mutants and transgenic strains used in the study are listed in Supplementary Data Table 1.

### Molecular biology and transgenic worms

All the constructs derived in this study were inserted into the pPD95.77 vector ^50^. For the *pezo-1*p1, p2, p3, and p4 promoter analysis, approximately 4 kb upstream sequences from each g, k, i, and l isoform start codon regions were amplified by PCR, respectively. To generate transgenic worms, 50 ng /μl of each reporter construct was injected, with 50 ng /μl of *unc-122*p::*dsRed* as an injection marker. For the *pezo-1* rescue experiments, a 4 kb fragment of *pezo-1*p1 promoter and a 1kb of *valv-1* promoter were fused with *pezo-1* cDNA and genomic DNA. 0.1 ng /μl of each construct was injected with 50 ng /μl of *unc-122*p::*dsRed* marker. For the Ca^2+^ imaging experiment, a 202 bp fragment of *ifa-4*Δ4 promoter was fused into a GCaMP6s vector ^51^. 100 ng/μl of construct was injected with 50 ng /μl of *unc-122*p::*dsRed* marker. To generate construct for *pezo-1* ectopic expression in the AWC neurons, a 568 bp fragment of *ceh-36*Δ1 promoter ^52^ was fused with *pezo-1* cDNA and 0.1 ng /μl of construct was injected with 50 ng /μl of *unc-122*p::*dsRed* marker.

### Microsphere movement assay

For microsphere movement assay, previous methods were modified^25^. Fluoresbrite Polystyrene YG microsphere was diluted with OP50 broth in a 1:9 ratio. 200 μl mixture was spread on a 6 cm NGM agar plate in a clean bench, which was placed at room temperature for 2 hr. Well-fed young adult animals were placed on a microsphere-contained plate for 20 min and transferred to an *E.coli* OP50 seeded plate. In 10 min after transferring worms, fluorescence time lapse images were acquired using an Axio cam 208 color with 6.3X objective and ZEN software for 10 min duration. Images obtained at 0 min, 5min, and 10 min were analyzed using Image J ^53^, and average fluorescence intensities were calculated by using GraphPad Prism.

### Pharyngeal plunge assay

Well-fed young adult worms were used to observe pharyngeal plunge under freely moving conditions. Each worm was transferred onto an *E.coli* OP50 spread-2% agarose pad on a glass slide and a coverslip was placed on top. Time lapse images were acquired over a 40 sec duration using a Zeiss Axio observer A1 with a 20X objective and Image J plugin software. At least 100 frames (20 sec) of continuous pharyngeal plunge movements from each worm were obtained and analyzed using Image J and GraphPad Prism.

### Hydropathy analysis

Transmembrane topology prediction and classification of PEZO-1 was performed using the DeepTMHMM (https://dtu.biolib.com/DeepTMHMM) program provided by the DTU-BioLib ^54^.

### Cell ablation by miniSOG

The cell ablation by miniSOG has been modified from previous methods ^29,30^. *valv-1*p::*miniSOG* transgenic animals were grown in foil-covered boxes until the experimental manipulation. A 460nm blue light source at approximately 2 mW/square mm as measured with an optical power/energy meter was originated from the Leica High-performance Fluorescence Stereomicroscope M205FA. Well-fed 15-20 young adult-stage *valv-1*p::*miniSOG* transgenic worms were transferred to a 3 cm NGM plate, and were exposed to the blue light in the dark for 1 hr. Animals were then recovered in the dark for 4 hr at a 15 ℃ incubator, and further experiments were conducted.

### *In vivo* calcium imaging in freely moving condition

Well-fed young adult-stage transgenic worms expressing GCaMP in the PI valve or head and neck muscle were used. Each worm was transferred onto a 3% agarose pad on a glass slide and covered with a coverslip. Fluorescence time-lapse images were acquired over a 40-second duration using a Zeiss Axio observer A1 with 40X or 20X objectives and analyzed using Image J. At least 100 frames (20 sec) during pharyngeal plunge from each worm were obtained and analyzed with a customized program that normalized the fluorescence intensity of the PI valve and muscles and subtracted the background.

### Serial Block Face Scanning Electron Microscopy (SBF-SEM)

The wild-type and *pezo-1* mutant animals were fixed with 2.5% glutaraldehyde in 0.1 M cacodylate buffer (pH 7.4) overnight at 4°C. After washing in CB, they were incubated in a reduced osmium solution (2% OsO4, 1.5% KCH in CB) for 1 hr, and the solution was replaced with filtered 1% TCH in D.W for 20 min. After washing in D.W., the solution was replaced with a 1% OsO4 aqueous solution for 30 min. Following washing, the solution was replaced with 1% UA in D.W. and incubated at 4°C overnight. The next day, the samples were washed with D.W. and incubated in a Lead aspartate solution at pH 5.5 for 30 min in a 50°C oven. After washing in D.W. six times for 10 min each, dehydration was performed through a graded ethanol series (30%, 50%, 70%, 95%, 100%, 30 min each at 4°C, pure ethanol 100% twice at room temperature). An ethanol: acetone mixture solution (1:1, 30 min, once), acetone (twice, 15 min each), acetone: Spurr’s (3:1, 1:1, 1:3, 1 hr each), and pure Spurr’s resin (overnight at room temperature) were sequentially used to infiltrate the resin. The next day, samples were incubated in fresh pure resin for 6 hr and placed in embedding molds in an oven (60°C) for 24 to 48 hr. Serial sections with a thickness of 100 nm were obtained from embedded samples on a plasma-treated silicon wafer for SEM imaging using an ultramicrotome (MT-X, RMC) equipped with ASH2 (RMC). EM data were obtained under 2 keV, 0.2 nA, 5 nm pixel size, 5 µs dwell time, 4096 X 4096 scan size, and a T1 backscattered electron detector using scanning electron microscopy (TeneoVS, FEI). The alignment, segmentation, and three-dimensional reconstruction were performed using the IMOD software^55^.

### Live imaging to assess the expression pattern of PEZO-1::mScarlet

For imaging the expression pattern of PEZO-1::mScarlet at the PI valve, the animals were immobilized on 7.5% agar pads with anesthetic buffer including 0.1% tricaine and 0.01% tetramisole, and M9 buffer. DIC and mScarlet image acquisitions were captured by a Nikon 60 X oil objective with 1 μm z-step size; 25-30 z planes were captured. A spinning disk confocal imaging system, including a Photometrics Prime 95B EMCCD camera and a Yokogawa CSU-X1 confocal scanner module was used to capture the images. The data were processed using Nikon NIS imaging software and ImageJ/FIJI Bio-formats plugin ^56^.

### Optogenetic experiments

L4 transgenic worm larvae expressing ReaChR::mKate2 transgenes under the control of indicated promoters were transferred 12 hr before the assay to either normal *E.coli* OP50 plates or *E.coli* OP50-retinal plates containing 1 mM all-trans-retinal (ATR, Sigma). *E.coli* OP50-retinal plates were prepared by seeding 200 μl *E.coli* OP50 with 2 μl of 100 mM ATR in 100% ethanol. To stimulate ReaChR, 565 nm LED at roughly 0.1 mW/square mm, as measured with an optical power/energy meter was used. At least 10 young adult hermaphrodites per strain were exposed in the presence or absence of ATR under a custom automated worm-tracking system. Time-lapse images were acquired over a 20-second duration using a Zeiss Axio observer A1 with a 40X objective and Image J plugin software. The heads of worms were allowed to move on the 3% agar pad, and recordings began with 5 sec in the absence of a green light, followed by 10 seconds of green light stimulation, and finally another 5 sec without a green light. The pharyngeal plunge was determined within 1 sec after supplying the green light stimulus.

### Inflating the anterior part of the intestine assay

For needle fabrication, Kwik-fill borosilicate capillary glass (1B100F-4) were pulled with Micropipette puller P-1000 (Detail setting of P-1000; Pull = 20, Vel. = 40, Delay = 20, Pressure = 130 at line 1×1 and Pull = 5, Vel. = 40, Delay = 110 at line 2×2). The injection solution (S-basal) was loaded onto the fabricated needle. Well-fed young adult worms were used for injection assay and fixed under a Zeiss Axio observer A1 with a 20X objective. The automatic microinjection FemtoJet was programmed for anterior intestinal injection (Detail setting of FemtoJet; injection pressure = 1700, injection time = 0.1 sec to 1 sec, compensation pressure = 102). NARISHIGE micromanipulator was used to hold the injection needle. The injection needle was oriented parallel to the anterior intestine plane and obliquely inserted.

### Confocal microscopy

Worms for microscopy were picked onto 2% agarose pads and anesthetized with 50 mM sodium azide in M9 buffer. At least 10 worms were imaged per genotype to obtain representative images. All images were acquired using a Zeiss LSM780 laser-scanning confocal. Z-stack images (each approximately 0.75 μm thick) were acquired using the Zen software and analyzed using the Zen software or Image J.

### Identification of neuronal expression pattern

Cell identification was performed by assessing position and size using Nomarski optics and by crossing with NeuroPAL (*otIs699*, *otIs670*)^57^. Imaging experiments using the NeuroPAL line were conducted using a Zeiss LSM780 laser-scanning confocal, with settings available for download on yeminilab.com. Analysis of NeuroPAL images for cell identification was conducted as described in Yemini et al. (2021) ^57^.

### Cross-correlation analysis

The JMP software (SAS) was used, as described, to analyze time series of PI valve movements with calcium fluorescence intensities, and head-bending angles with calcium fluorescence intensities for a cross-correlation analysis ^58^. The time lag was 10 sec, and PI valve position and head position was used as an input. Peak correlation represents the correlation values at lag 0 sec and is compared using the ANOVA test.

### Quantification and Statistical analysis

GraphPad Prism software was used for data analysis. Comparisons and P-value calculations were made between animals of the same or different strains, and treated and untreated animals, using Student’s t-test, one-way ANOVA, and two-way ANOVA with corrections for multiple hypothesis testing. More statistical information is represented in all figure legends.

## Acknowledgments

We are grateful to the *Caenorhabditis* Genetics Center (NIH Office of Research Infrastructure Programs, P40 OD010440) and the National BioResource Project (Japan) for strains. We are also grateful to Andy Golden, Piali Sengupta, Chris Li, and K. Kim Labs for helpful discussion and/or critical comments on this manuscript and Sujin Jo and Taehyun Kim for their technical assistance. This work was supported by the National Research Foundation of Korea (NRF-2021R1A2C1008418, NRF-2020R1A4A10) (K.Kim), and the National Research Foundation of Korea (2021R1A2B5B01002702) (K.Kang).

## Author contributions

Y.P., J.Y., B.X., K.Ka., K.L., and K.Ki. conceived and designed the study. Y.P, J.Y., J.C., D.K., B.X., Y. O., J.K., H.N., H. H., W.H., J.K. and S.J. performed the experiments. Y.P., J.Y., K.L. and K.Ki. analyzed the data. Y.P., J.Y. and K.Ki. wrote the paper. All authors read, edited and approved the final manuscript.

