## Supplementary figures for "PIEZO acts in an intestinal valve to regulate swallowing in *C. elegans*"

### Supplementary Fig. 1

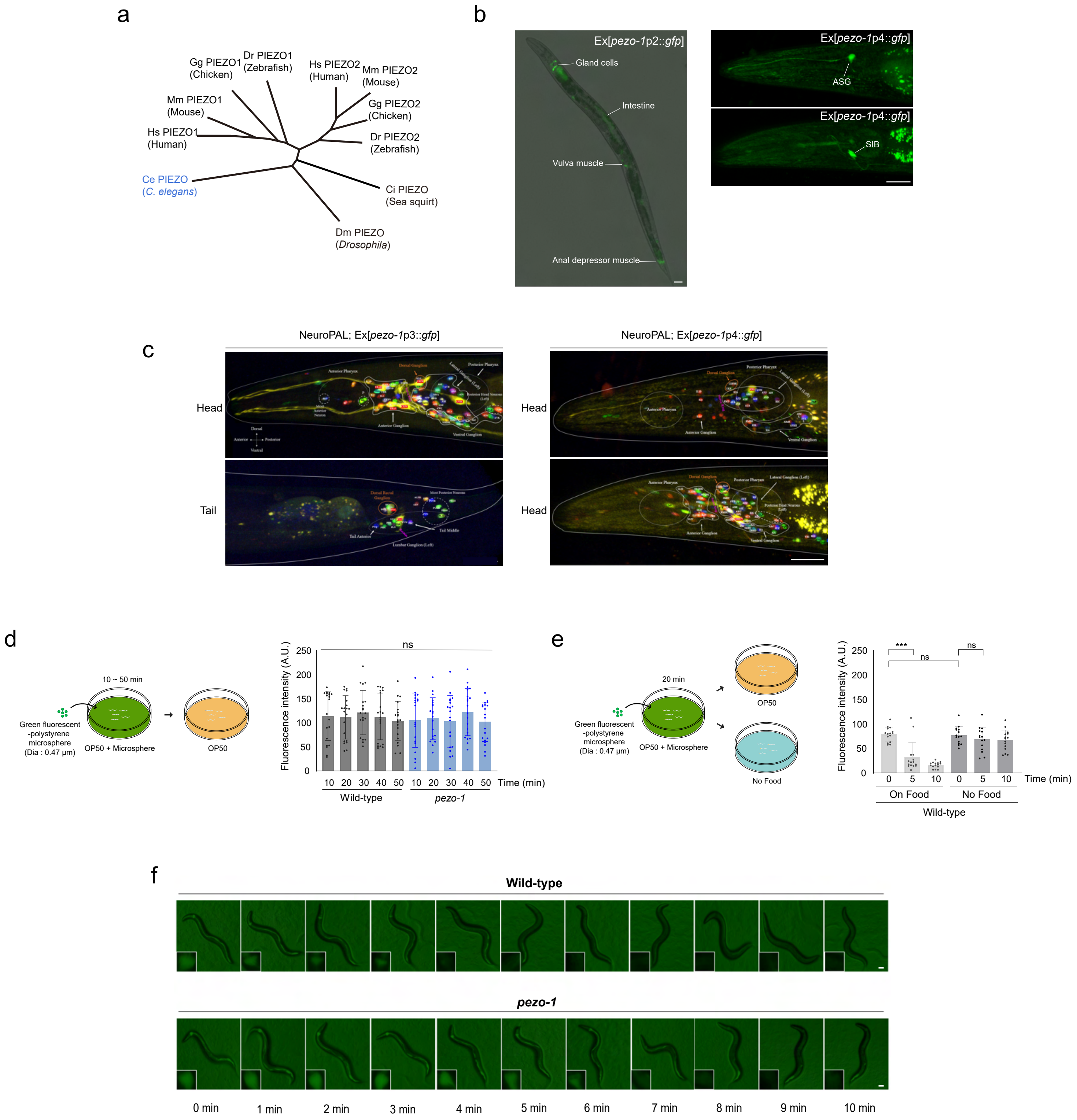

**Supplementary Fig. 1: PEZO-1 regulates food movement in the intestine.**

(a) Phylogenetic tree of the PIEZO family. Hs, *Homo sapiens*; Mm, *Mus musculus*; Gg, *Gallus gallus*; Dr, *Danio rerio*; Ci, *Ciona intestinalis*; Dm, *Drosophila melanogaster*; Ce, *Caenorhabditis elegans*. (b) Representative images of wild-type animals expressing *pezo-1*p2::*gfp* (left) and *pezo-1*p4::*gfp* (right) transgene. Anterior is to the left. Scale bar: 10 μm. (c) Neuronal cell identification using NeuroPAL worm. The merge images of transgenic animals co-expressing NeuroPAL and *pezo-1*p3::*gfp* and *pezo-1*p4::*gfp* (yellow) in the head and tail. (d-e) Experimental scheme (left) and fluorescence intensity of microspheres in the anterior part of the intestine (right) of indicated worms according to changes in microsphere exposure time (d) and presence or absence of OP50 (e). Each dot represents the fluorescence intensity of a single animal. n=15 each. Error bars indicate SD. *** indicates significant differences at $p<0.001$ (one-way ANOVA test followed by the Tukey test). (f) Images of the whole body and anterior part of the intestine (boxed regions) of wild-type and *pezo-1* (*tm10725*) mutants taken every minute for 10 minutes after recovery from microsphere ingestion. Scale bar: 50 μm.

### Supplementary Fig. 2

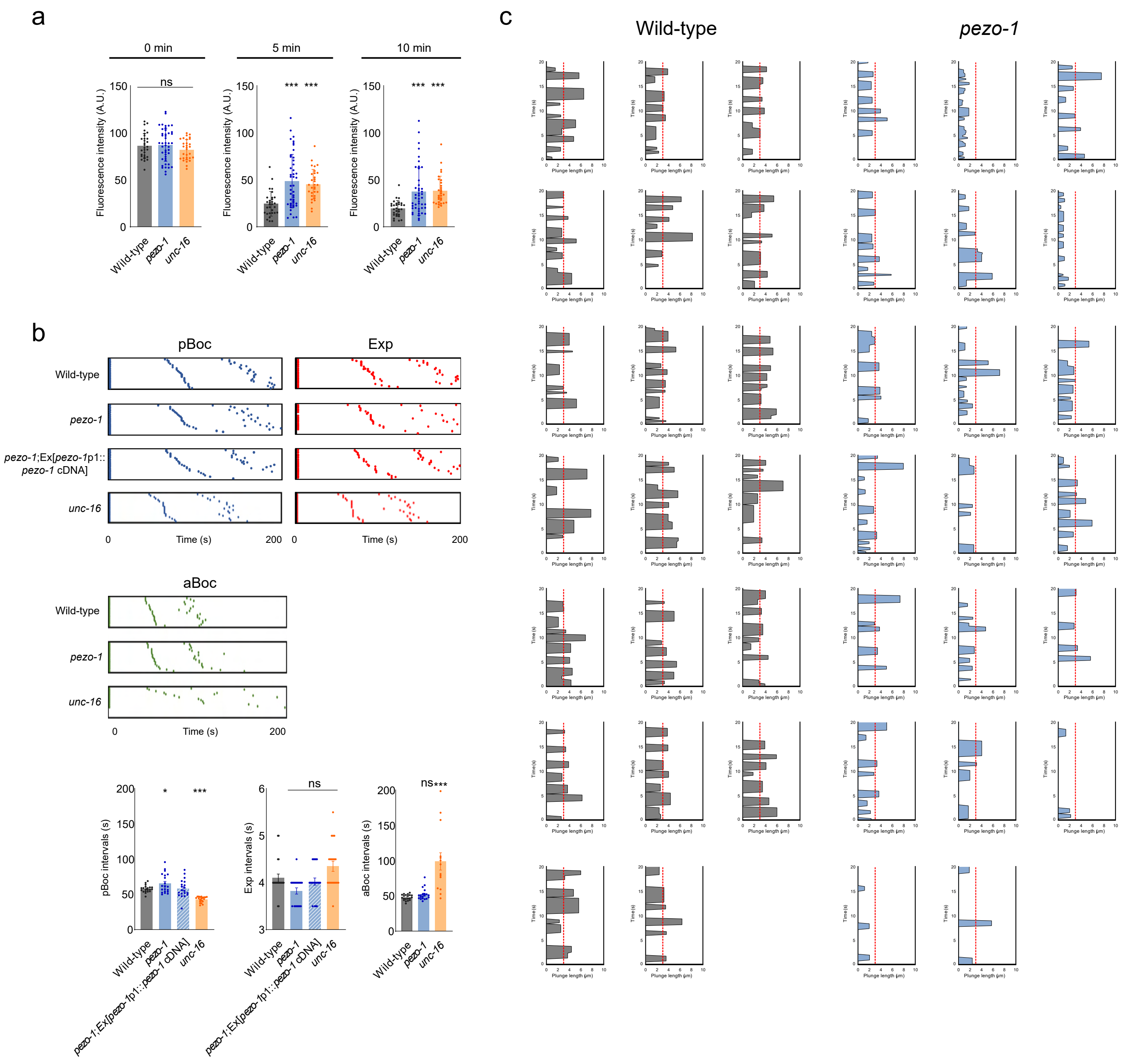

**Supplementary Fig. 2: *pezo-1* mutants exhibit defects in pharyngeal plunge.**

(a) Fluorescence intensity of microsphere in the anterior part of the intestine of wild-type animals and *pezo-1* (*tm10725*) or *unc-16* (*e109*) mutants. n ≥ 30 for each genotype. Error bars indicate SD. *** indicates significant differences from wild-type at $p<0.001$ (one-way ANOVA test followed by the Dunnett test). (b) Defecation cycle of the indicated genotype. Blue, red, and green dots indicate the time when pBoc (posterior body muscle contraction), Exp (Expulsion), and aBoc (anterior body muscle contraction) are observed after the first event happens (time=0) (upper). n=20. Note that five *unc-16* mutants did not exhibit aBoc. The average interval times for pBoc, Exp and aBoc (below). n=20. Error bars indicate SEM. * and *** indicate significant differences from wild-type at $p<0.05$ and $p<0.001$, respectively (one-way ANOVA test followed by the Dunnett test). The time series plot of plunge length in wild-type (left) and *pezo-1* (*tm10725*) mutant (right) animals during a 20-second pharyngeal plunge. (c) The time series plot of plunge length in wild-type (left) and *pezo-1* (*tm10725*) mutant (right) animals during a 20-second pharyngeal plunge.

#### Supplementary Fig. 3

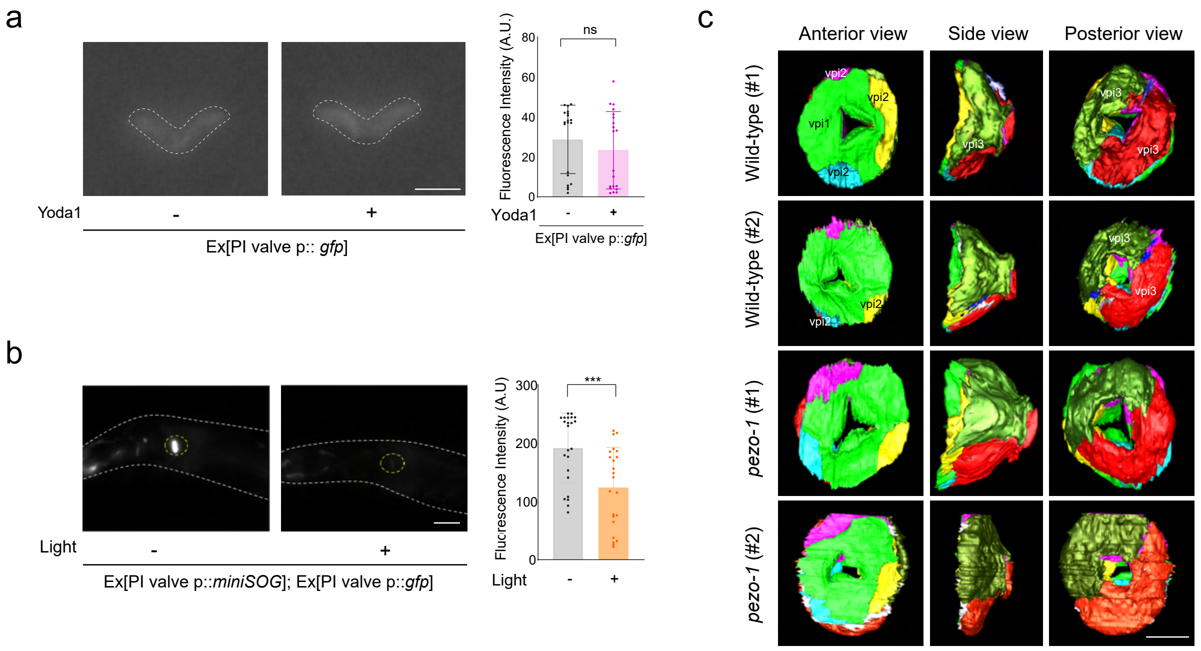


**Supplementary Fig. 3: The PI valve mediates pharyngeal plunge.**

(a) Representative fluorescence images of PI valve in Yoda1-exposed animals (left). Scale bar: 5 µm. The average fluorescence intensity of PI valve in Yoda1- exposed animals (right). n = 20. Error bars indicate SD. ns indicates no significant differences from Yoda1-unexposed wild-type (unpaired t-test). (b) Representative fluorescence images (left) and the average fluorescence intensity (right) of the PI valve in a genetically ablated animal co-expressing *miniSOG* and *gfp* genes under the control of PI valve-specific *valv-1* promoter. Scale bar: 10 µm. n ≥ 20. Error bars indicate SD. *** indicates significant differences at $p<0.001$ (unpaired t-test). (c) The Serial Block Face Scanning Electron Microscopy images of PI valve in the anterior view (left), side view (middle) and posterior view (right), respectively. Each of PI valve cells is labeled in green (vpi3), red (vpi3), yellow (vpi2), magenta (vpi2), cyan (vpi2) and light green (vpi1) respectively. Scale bar: 5 µm.

### Supplementary Fig. 4

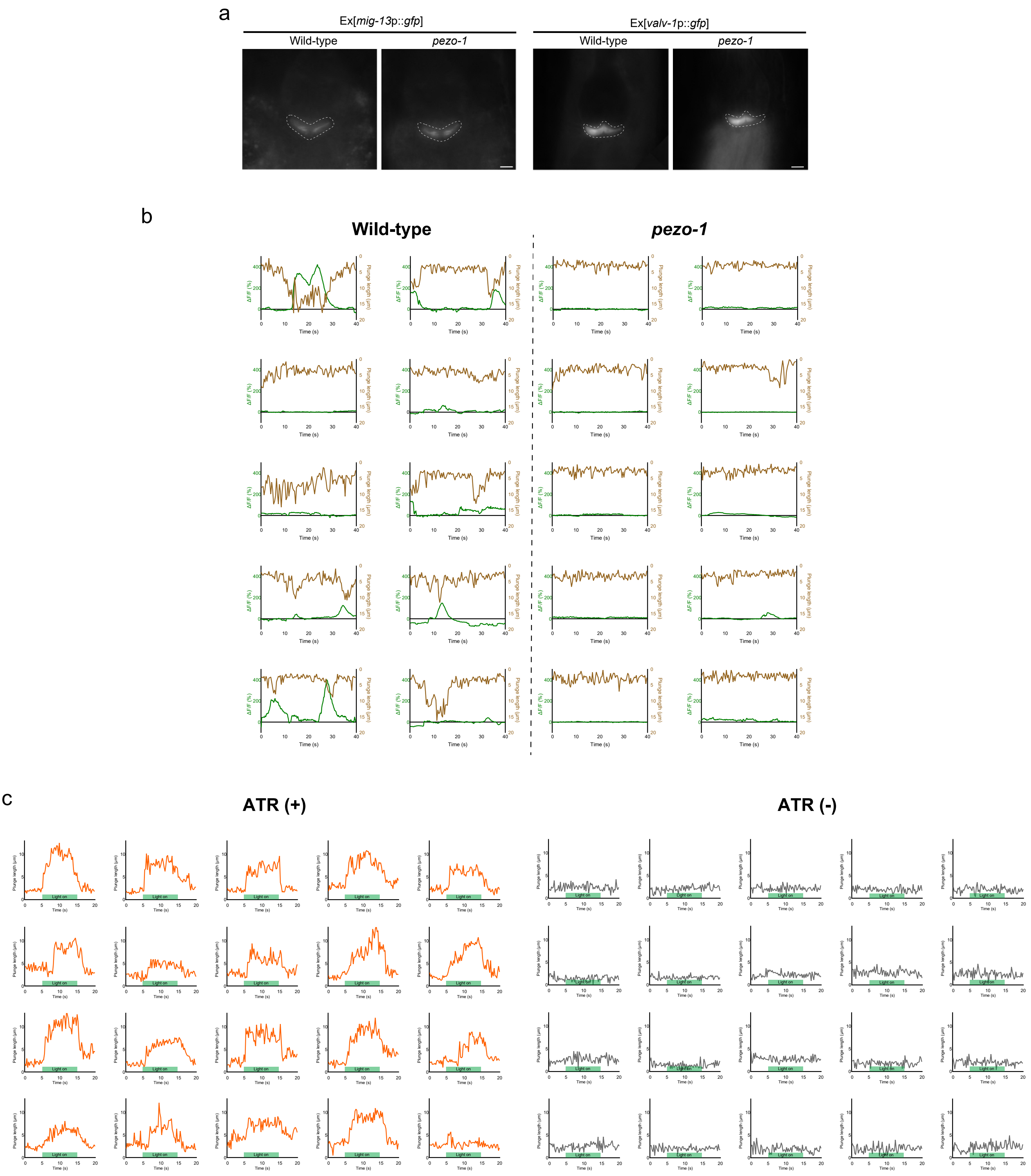

**Supplementary Fig. 4: $Ca^{2+}$ activaties in the PI valve mediate pharyngeal plunge.**

(a) The representative PI valve fluorescence images under the control of *mig-13* promoter (left) and *valv-1* promoter in *pezo-1* (*tm10725*) mutants. Scale bar: 5 µm. (b) Calcium dynamics (green) in the PI valve and the corresponding plunge length (brown) in the same wild-type or *pezo-1* (*tm10725*) mutant animals. (c) Plunge length after 10 seconds of light stimulation in the presence or the absence of all trans retinal (ATR).

### Supplementary Fig. 5

a

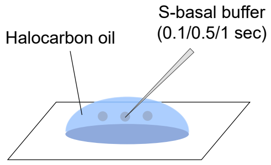


| | 0.1 sec | 0.5 sec | 1 sec |
|---|---|---|---|
| Average r (µm) | 6.455 | 9.155 | 21.69 |
| $4/3r^3\pi$ ($\mu m^3$) | 1126.049 | 3212.502 | 42743.2 |
| Volume (pl) | 2.2 | 6.3 | 42.7 |

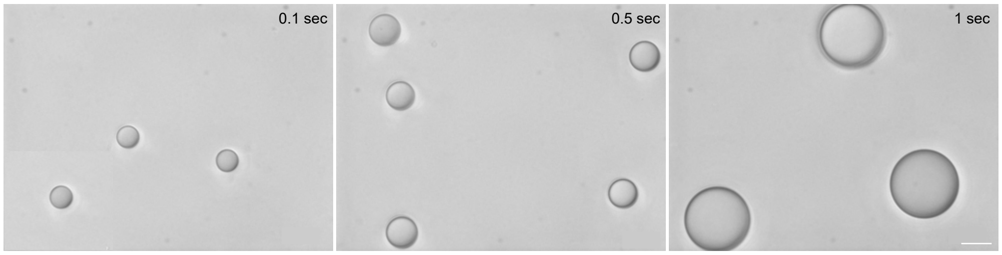


b

**Control**

**Ex[AWCp ::*pezo-1* cDNA]**

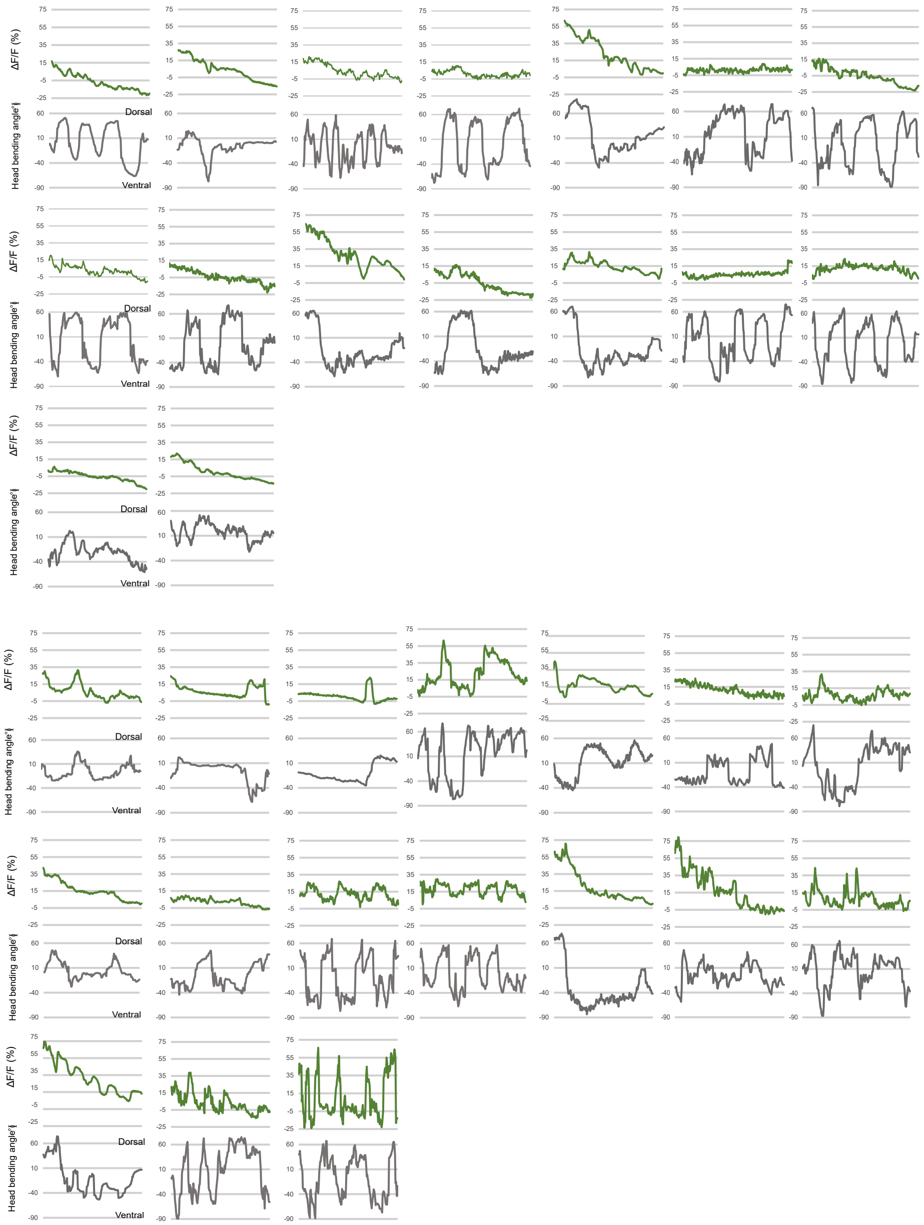

**Supplementary Fig. 5: PEZO-1 detects mechanical forces.**

(a) Schematic of how to calculate the volume of injected solution. The injected S-basal buffer forms a perfect sphere in halocarbon oil. Representative images of buffer spheres in halocarbon oil using the same injection needle are shown. The radius was measured for each sphere after injection time of 0.1, 0.5, and 1 second. (b) Traces of AWC GCaMP activities (upper) and the corresponding head bending angle (lower) of control (top) or AWCp::*pezo-1* cDNA transgenic animals (bottom).

**Supplementary Fig. 6**

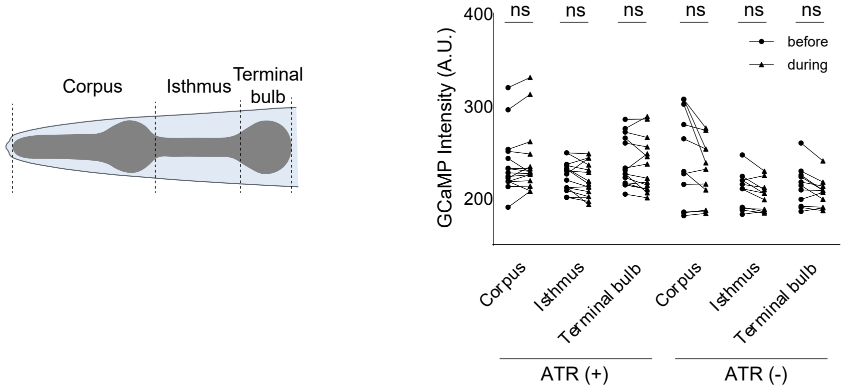


**Supplementary Fig. 6: The pharyngeal plunge is not mediated by pharyngeal muscle contraction.**

Schematic of pharyngeal muscles in *C. elegans* (left). Calcium activities in corpus, isthmus, and terminal bulb of transgenic animals expressing GCaMP3 in the pharyngeal muscles and ReaChR in the PI valve before and during the plunge. The pharyngeal plunge was elicited by light stimulation in the presence or the absence of all trans retinal (ATR). n ≥ 10. ns indicates no significant differences (two-way ANOVA test followed by the Sidak's test).

### Supplementary Data Table. 1

| Genotype | Strain | |
|---|---|---|
| *lskEx1214*[*pezo-1*p1::*gfp, unc-122*p::*dsRed*] | KHK1757 | |
| *lskEx901*[*pezo-1*p3::*gfp, unc-122*p::*dsRed*] | KHK1292 | |
| *pezo-1 (tm10725)* | KHK1839 | |
| *pezo-1 (tm10726)* | - | |
| *pezo-1 (tm5071)* | - | |
| *pezo-1 (tm5750)* | - | |
| *pezo-1 (tm5111)* | - | |
| *pezo-1 (lsk51)* | KHK1771 | |
| *pezo-1(tm10725)* ; *lskEx1294*[*pezo-1*p1::*pezo-1* g isoform cDNA, *unc-122*p::*dsRed*] | KHK1848 | |
| *pezo-1(tm10725)* ; *lskEx1298*[*pezo-1*p3::*pezo-1* i isoform cDNA, *unc-122*p::*dsRed*] | KHK1852 | |
| *pezo-1(tm10725)* ; *lskEx1305*[*pezo-1*p1::*pezo-1* i isoform cDNA, *unc-122*p::*dsRed*] | KHK1859 | |
| *pezo-1(tm10725)* ; *lskEx1176*[*pezo-1*p3::*pezo-1* g isoform cDNA, *unc-122*p::*dsRed* ] | KHK1711 | |
| *pezo-1(tm10725)* ; *lskEx1697*[*valv-1*p::*pezo-1* g isoform cDNA, *unc-122*p::*dsRed*] | KHK2381 | |
| *pezo-1(tm10725)* ; *lskEx1604*[*acd-5*p::*pezo-1* g isoform cDNA, *unc-122*p::*dsRed*] | KHK2250 | |
| *pezo-1(tm10725)* ; *lskEx1617*[*myo-2*p::*pezo-1* g isoform cDNA, *unc-122*p::*dsRed*] | KHK2277 | |
| *pezo-1(tm10725)* ; *lskEx1314*[*pezo-1*p1::mPiezo1 cDNA, *unc-122*p::*dsRed*] | KHK1868 | |
| *unc-16 (e109)* | CB109 | |
| *lskEx1351*[*valv-1*p::miniSOG, *unc-122*p::*dsRed*] | KHK1906 | |
| *lskEx1683*[*ifa-4Δ4*p::GCaMP6s, *unc-122*p::*dsRed*] | KHK2364 | |
| *pezo-1(tm10725)* ; *lskEx1683*[*ifa-4Δ4*p::GCaMP6s, *unc-122*p::*dsRed*] | KHK2444 | |
| *lite-1* ; *lskEx1337*[*myo-2*p::GCaMP3; *valv-1*p::ReaChR::mKate2, *unc-122*p::*dsRed*] | KHK1916 | |
| *av182*[*pezo-1*::mScarlet] | AG483 | |
| *lskEx1553*[*ceh-36Δ1*p::GCaMP3, *unc-122*p::*dsRed*] | KHK2169 | |
| *lskEx1553*[*ceh-36Δ1*p::GCaMP3, *unc-122*p::*dsRed*]; *lskEx1581*[*ceh-36Δ1*p::*pezo-1* g isoform cDNA, *unc-122*p::*gfp*] | KHK2208 | |
| *goeIs3*[*myo-3*p::GCaMP3.35] | HBR4 | |
| *unc-31 (e928)* | CB928 | |
| *unc-13 (e1091)* | CB1091 | |
| *tdc-1 (n3419)* | MT13113 | |
| *unc-49 (tm5487)* | - | |
| *tbh-1 (n3247)* | MT9455 | |
| *tph-1 (n4622)* | MT14984 | |
| *unc-17 (e245)* | CB933 | |
| *cat-2 (e1112)* | CB1112 | |
| *eat-4 (ky5)* | MT6308 | |
| *lskEx1493*[*pezo-1*p2::*gfp, unc-122*p::*dsRed*] | KHK2085 | |
| *lskEx1702*[*pezo-1*p4::*gfp* , *unc-122*p::*dsRed*] | KHK2390 | |
| NeuroPAL *(otIs699)* ; *lskEx901[pezo-1p3::gfp, unc-122p::dsRed]* | KHK2491 | |
| NeuroPAL *(otIs670)* ; *lskEx1702*[*pezo-1*p4::*gfp* , *unc-122*p::*dsRed*] | KHK2492 | |
| *pezo-1*(*tm10725*); *lskEx1718*[*pezo-1*p1::mPiezo2, *unc-122*p::*dsRed*] | KHK2414 | |
| *pezo-1*(*tm10725*); *lskEx1738*[*pezo-1*p1::mPiezo2, *unc-122*p::*dsRed*] | KHK2445 | |
| *lskEx1679*[*ifa-4Δ4*p::*gfp,unc-122p*::*dsRed*] | KHK2358 | |
| *lskEx1351*[*valv-1*p::miniSOG, *unc-122*p::*dsRed*]; *lskEx1224*[*valv-1*p::*gfp, unc-122*p::*dsRed*] | KHK1980 | |
| *lskEx1479*[*mig-13*p::*gfp, unc-122p*::*dsRed*] | KHK2063 | |
| *pezo-1(tm10725); lskEx1479*[*mig-13*p::*gfp, unc-122p*::*dsRed*] | KHK2064 | |
| *lskEx1224*[*valv-1*p::*gfp, unc-*122*p*::*ds* Red] | KHK1772 | |
| *pezo-1(tm10725)* ; *lskEx1224*[*valv-1*p::*gfp, unc-122*p::*dsRed*] | KHK2379 | |
| **Chemicals** | | |
| Yoda1 | Sigma-Aldrich | Cat#448947-81-7 |
| Fluoresbrite Polystyrene YG microsphere | Polysciences | Cat#17152-10 |
| 5-fluoro-2'-deoxyuridine | Sigma-Aldrich | Cat#50-91-9 |
| **Commercial assays** | | |
| iProof High-Fidelity DNA Polymerase | BIO-RAD | Cat#1725301 |
| QIAquick PCR & Gel Cleanup Kit (100) | QIAGEN | Cat#28506 |
| KOD FX Neo-High fidelity DNA polymerase | TOYOBO | Cat#045700 |
| Plasmid DNA Miniprep S&V Kit | BIONICS | Cat#DN10200 |

**Supplementary Video 1 and 2.** Pharyngeal plunge in wild-type animal and *pezo-1* mutant. The animals were recorded for 20 seconds. The white circle and vertical line indicate the terminal bulb of the pharynx and the anterior end of the intestine, respectively. The red dots mark the occurrence of the pharyngeal plunge.

**Supplementary Video 3 and 4.** Calcium dynamics in the PI valve. The wild-type animal and *pezo-1* mutant expressing GCaMP6 in the PI valve was placed on a 3% agar pad and recorded for 20 seconds. The white circle and vertical line indicate the terminal bulb of the pharynx and the anterior end of the intestine, respectively.

**Supplementary Video 5 and 6.** Optogenetic activation of the PI valve in transgenic animals in the presence or absence of all trans retinal (ATR). The white circle and vertical line indicate the terminal bulb and the anterior end of the intestine, respectively. The red dots mark the occurrence of the pharyngeal plunge.

**Supplementary Video 7 and 8.** Worm's head during injecting buffer in wild-type animal and *pezo-1* mutant. Approximately 50 pl buffer was microinjected. The white circle and vertical line indicate the terminal bulb and the anterior end of the intestine, respectively. The red dots mark the occurrence of the pharyngeal plunge.
